# An oomycete effector co-opts a host RabGAP protein to remodel pathogen interface and subvert defense-related secretion

**DOI:** 10.1101/2024.01.11.575225

**Authors:** Enoch Lok Him Yuen, Yasin Tumtas, Lok I Chan, Tarhan Ibrahim, Edouard Evangelisti, Frej Tulin, Jan Skłenar, Frank Menke, Sophien Kamoun, Doryen Bubeck, Sebastian Schornack, Tolga O. Bozkurt

## Abstract

Pathogens have evolved sophisticated mechanisms to manipulate host cell membrane dynamics, a crucial adaptation to survive in hostile environments shaped by innate immune responses. Plant-derived membrane interfaces, engulfing invasive hyphal projections of fungal and oomycete pathogens, are prominent junctures dictating infection outcomes. Understanding how pathogens transform these host-pathogen interfaces to their advantage remains a key biological question. Here, we identified a conserved effector, secreted by plant pathogenic oomycetes, that co-opts a host Rab GTPase-activating protein (RabGAP), TBC1D15L, to remodel the host-pathogen interface. The effector, PiE354, hijacks TBC1D15L as a susceptibility factor to usurp its GAP activity on Rab8a—a key Rab GTPase crucial for defense-related secretion. By hijacking TBC1D15L, PiE354 purges Rab8a from the plasma membrane, diverting Rab8a-mediated immune trafficking away from the pathogen interface. This mechanism signifies an uncanny evolutionary adaptation of a pathogen effector in co-opting a host regulatory component to subvert defense-related secretion, thereby providing unprecedented mechanistic insights into the reprogramming of host membrane dynamics by pathogens.

## Introduction

Plants are equipped with a dynamic innate immune system to sense and confront pathogens. This system fundamentally relies on endomembrane trafficking, which facilitates a hostile environment against pathogens by directing immune components, such as pathogenesis-related (PR) proteins, to the pathogen interface. Consistent with this notion, a growing number of studies have revealed pathogen manipulation of plant vesicle trafficking as a ubiquitous infection strategy^1–6^.

Pathogens intimately interact with plant cells via specialized structures that facilitate the transfer of effector proteins and the uptake of nutrients. Filamentous plant pathogens, including oomycetes and fungi, project hyphal extensions that breach the cell wall and penetrate host cells. At these junctures, plants mount targeted immune responses, which include cellular reinforcements and secretion of defense molecules^7–10^. Filamentous pathogens have developed strategies to overcome these defenses, forming specialized infection structures like haustoria or infection vesicles (formed by oomycete pathogens), which are accommodated inside host cells. These infection structures are enveloped by plant-derived membranes with unique biochemical compositions, often lacking transmembrane proteins including pattern recognition receptors, delineating a polarized membrane interface^11,12^. At these interfaces, pathogens are thought to manipulate the environment, creating safe niches for efficient effector delivery and nutrient absorption. However, the regulatory mechanisms governing the trafficking of immune components at the host-pathogen interface and the extent to which they are manipulated by pathogen effectors remain largely unknown.

Rab GTPases (Rabs) are integral to vesicle trafficking and immunity, mediating the movement and fusion of vesicles with membrane compartments^13^. While the immune functions of plant Rabs remain largely unknown, members of the Rab8 and Rab11 have been identified to contribute to pathogen resistance by facilitating defense-related secretion^3,4,14^. Rabs function as molecular switches cycling between GTP-bound active and GDP-bound inactive states. Their activation is regulated by guanine nucleotide exchange factors (GEFs), which facilitate GTP loading, and their inactivation is mediated by GTPase-activating proteins (GAPs), which accelerate GTP hydrolysis. Most RabGAPs are characterized by the Tre2/Bub2/Cdc16 (TBC) domain featuring dual catalytic fingers accelerating the GTP hydrolysis of their cognate Rabs^15^, thereby controlling their localization and functions. Although a few RabGAPs have been implicated in immunity^16,17^, the mechanisms behind their action, their specific Rab substrates, and the trafficking pathways they regulate in immune responses remain largely unexplored in both plants and animals.

The critical role of membrane trafficking in plant pathogen defense is increasingly evident, with diverse pathogens deploying effectors that virtually target every facet of vesicle trafficking^5,6,10,18–20^. Proteomic screens have highlighted the strategic targeting of the host vesicle trafficking system by *Phytophthora* pathogens^6,21^. Notably, these pathogens deploy effectors converging on key Rab GTPases, like Rab8 and Rab11, which are integral to defense-related secretion^3,4,14^. However, the detailed mechanisms of these interactions and their impact on host membrane dynamics are still not fully understood. Despite extensive documentation of effectors targeting host Rab GTPases in plant and animal pathosystems, the potential targeting of RabGAPs by pathogen effectors remains an unexplored area. This is particularly intriguing given the crucial role of RabGAPs in regulating Rab functions.

Here, we elucidate an unprecedented mechanism employed by a conserved effector family from the *Phytophthora* species, notably PiE354 and its homologs, in reconfiguring host cell membrane dynamics at the pathogen interface. PiE354 adeptly co-opts the host RabGAP protein TBC1D15L to harness its GAP activity on Rab8a. This manipulation expels Rab8a from the plasma membrane, redirecting Rab8a-mediated secretion of antimicrobial compounds away from the site of pathogen attack. Our findings suggest a detailed mechanistic model where PiE354 physically perturbs the TBC1D15L-Rab8a complex, leveraging the GAP activity of TBC1D15L to subvert Rab8a-mediated immune trafficking. Our research uncovers a sophisticated strategy employed by pathogen effectors, demonstrating how they exploit the catalytic functions of a host transport regulator to effectively remodel host membrane dynamics to subvert immune responses.

## Results

### A plant RabGAP protein is targeted by a conserved *Phytophthora* effector

To elucidate how *Phytophthora* species manipulate vesicle trafficking pathways, we focused on identifying effectors that target the plant endomembrane transport machinery. As part of an unbiased yeast two-hybrid (Y2H) screen, we discovered that the *P. palmivora* effector TIKI (trafficking interference and tissue killing effector, PLTG_0964243, Table S1) associates with a *Nicotiana benthamiana* TBC-containing RabGAP protein (Nbe.v1.s00100g29830, TBC1D15L hereafter) akin to members of the mammalian TBC1D15 family (Table S2). To corroborate the Y2H results, we conducted an immunoprecipitation-mass spectrometry (IP-MS) analysis, which again pinpointed TBC1D15L as a candidate interactor of TIKI (Table S3).

We validated the association between TIKI and TBC1D15L through *in planta* reverse co-immunoprecipitation (co-IP), pulling down N-terminally green fluorescent protein (GFP)-tagged TBC1D15L with a plant expression construct of TIKI effector domain N-terminally-tagged with red fluorescent protein (RFP) (Figure 1A). In *N. benthamiana* expression assays, we observed that TIKI triggers plant cell death (Figure S1A), presenting challenges for conducting accurate biochemical and cellular biology assays. However, the two TIKI homologs from *P. infestans* (Figure S1B), named PiE354 (*Phytophthora infestans* effector 354, PITG_04354, Table S1) and PiE355 (*Phytophthora infestans* effector 355, PITG_04355, Table S1), showed varying cell death responses. PiE355 induced less severe cell death than TIKI, whereas PiE354 showed no visible symptoms (Figure S1C), presenting an excellent opportunity for elucidating the functions of these effectors. AF2 structural predictions showed notable similarity among these three *Phytophthora* effectors, indicated by a low root-mean-square deviation (RMSD) value of 0.810 and 0.374 when comparing TIKI to PiE354 and PiE355, respectively, hinting at a conserved mode of action (Figures 1B and S1D). Consistent with this notion, co-IP experiments using protein extracts from *N. benthamiana* demonstrate that GFP:TBC1D15L interacts with both RFP:PiE354 and RFP:PiE355, but not with RFP:EV control (Figure 1C). Conversely, the GFP:EV control did not interact with any of the effectors (Figure 1C). Consistent with these findings, confocal microscopy analysis revealed that TBC1D15L colocalizes with all three effectors (TIKI, PiE354, and PiE355) at discrete punctate structures and in the cytosol. (Figure 1D). It is worth noting that while PiE354 exhibited puncta formation, the frequency was less than the other two effectors (Figure S1E). Altogether, these results show that TIKI and its *P. infestans* homologs, PiE354 and PiE355, target TBC1D15L in host plants.

**Figure 1.**
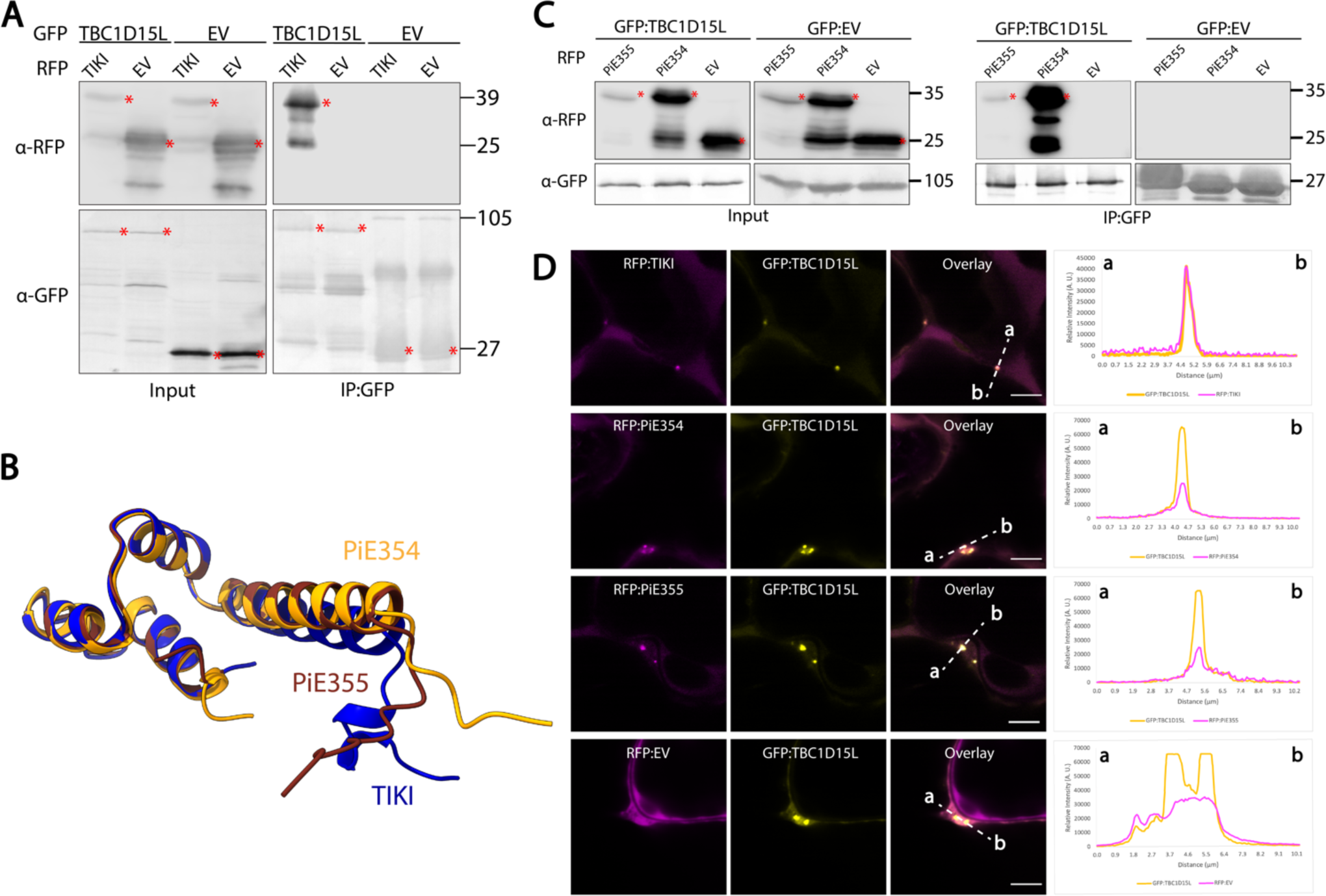
Conserved effectors from *Phytophthora* species target TBC1D15L. (A) TIKI interacts with TBC1D15L *in planta*. RFP:TIKI, or RFP:EV was transiently co-expressed with either GFP:TBC1D15L, or GFP:EV. IPs were obtained with anti-GFP antibody. Total protein extracts were immunoblotted. Red asterisks indicate expected band sizes. Numbers on the right indicate kDa values. (B) Structural alignment of the effectors TIKI (blue) from *P. palmivora*, PiE354 (orange) and PiE355 (brown) from *P. infestans*. Structural predictions were obtained via AF2. The model shows overall structural conservation of the effectors. (C) PiE355 and PiE354 interact with TBC1D15L *in planta*. GFP:TBC1D15L was transiently co-expressed with either RFP:PiE355, RFP:PiE354, or RFP:EV. IPs were obtained with anti-GFP antibody. Total protein extracts were immunoblotted. Red asterisks indicate expected band sizes. Numbers on the right indicate kDa values. (D) TBC1D15L colocalizes with TIKI, PiE354 and PiE355 in puncta *in planta*. Confocal micrographs of *N. benthamiana* leaf epidermal cells transiently expressing either RFP:TIKI (1^st^ row), RFP:PiE354 (2^nd^ row), RFP:PiE355 (3^rd^ row), or RFP:EV (4^th^ row), with GFP:TBC1D15L. Presented images are single plane images. Transects in overlay panels correspond to line intensity plots depicting the relative fluorescence across the marked distance. Scale bars, 5 µm.

### PiE354 targets the N-terminal Rab-binding domain of TBC1D15L

We next investigated the mechanisms by which PiE354 interacts with TBC1D15L. We chose PiE354 because, unlike its homologs TIKI and PiE355, it does not induce cell death in plants, thus avoiding complications in physiological and functional analyses (Figure S1C). Taking advantage of AF2, we first visualized the protein architecture of TBCD15L. This analysis revealed a domain of unknown function, DUF3548, at the N-terminus, and a TBC domain (TBCD) near the C-terminus (Figure S2A). The DUF3548, although not fully characterized, has been reported to function as a Rab-binding domain (RBD) in the human TBC-RabGAP protein RUTBC2^22^. Notably, AF2-multimer (AF2-M) structural predictions of the TBC1D15L-PiE354 complex indicated that PiE354 establishes multiple high-confidence contacts with the candidate RBD (DUF3548) and a few low confidence contacts with the TBCD, spanning a distance of about 5 Å (Figures 2A and S2B), suggesting that the effector targets the RBD of TBC1D15L. This finding aligns with our Y2H results (Table S2), which indicated that the N-terminal region of TBC1D15L is sufficient for binding TIKI.

**Figure 2.**
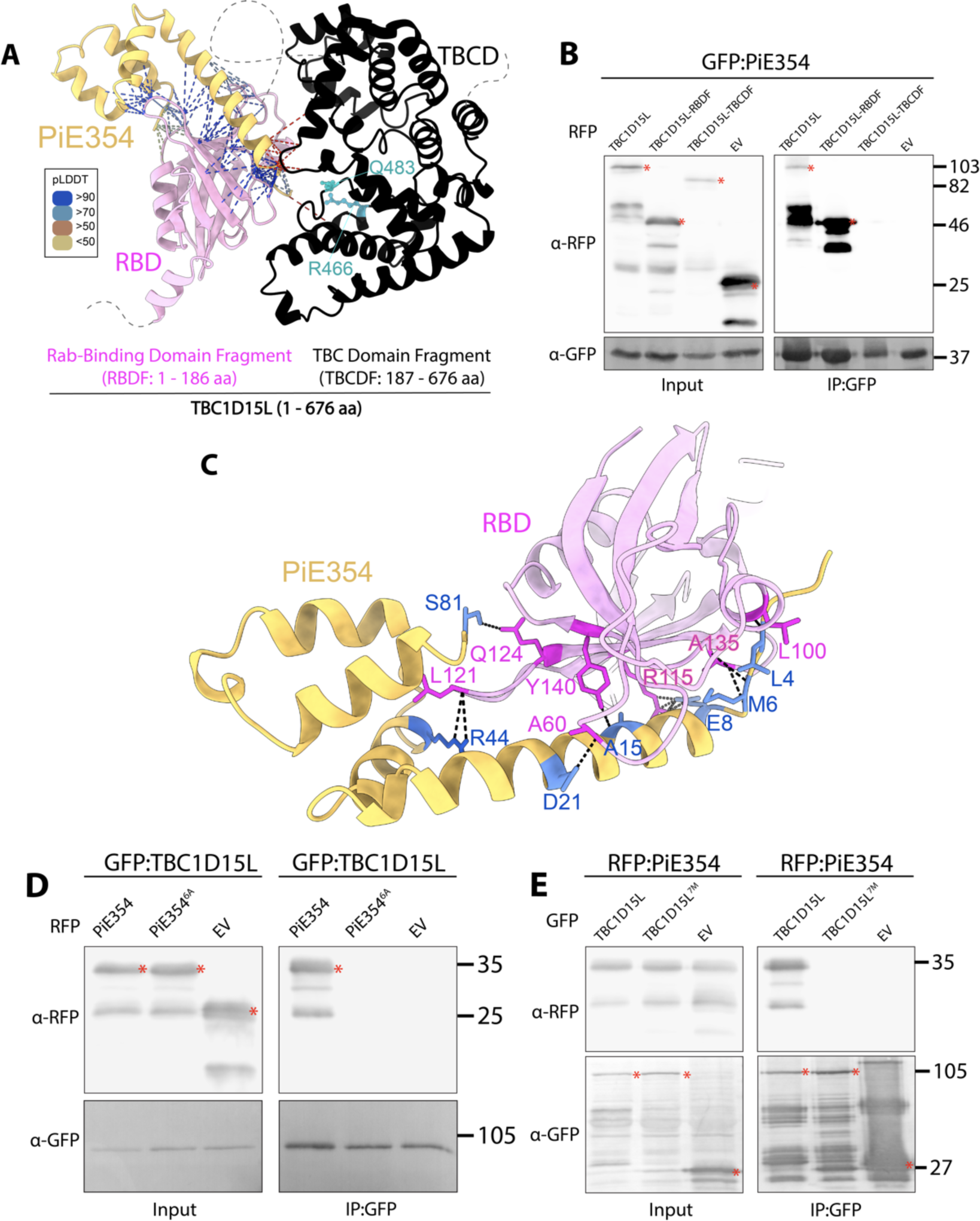
PiE354 targets the N-terminal RBD fragment of TBC1D15L. (A) AF2-M-predicted model of PiE354 targeting TBC1D15L. PiE354, RBD of TBC1D15L, and TBCD of TBC1D15L are depicted in yellow, pink, and black, respectively. The key residues responsible for the GAP activity of TBC1D15L, R466 and Q483, are highlighted in cyan. The bonds between PiE354 and TBC1D15L are predicted by ChimeraX with a distance of 5 Å. The colors of the bonds are based on the AF2-calculated prediction confidence score (pLDDT) as indicated in the rectangular box. (B) PiE354 interacts with full length TBC1D15L and RBDF of TBC1D15L, but not with TBCDF of TBC1D15L and EV. GFP:PiE354 was transiently co-expressed with either RFP:TBC1D15L, RFP:RBDF, RFP:TBCDF, or RFP:EV. (C) AF2-M-predicted model of PiE354 (yellow) targeting the RBD of TBC1D15L (pink), depicting their interacting residues. The bonds between PiE354 and TBC1D15L are predicted by PyMOL with a distance of 3 Å. The PiE354-RBD interaction interface consists of 7 key residues on both proteins, colored as blue on PiE354, and as magenta on RBD. (D) PiE354 targets TBC1D15L through 6 key residues on PiE354. GFP:TBC1D15L was transiently co-expressed with either RFP:PiE354, RFP:PiE354^6A^, or RFP:EV. (E) TBC1D15L interacts with PiE354 through 7 key residues on TBC1D15L. RFP:PiE354 was transiently co-expressed with either GFP:TBC1D15L, GFP:TBC1D15L^7M^, or GFP:EV. For all co-IP assays, IPs were obtained with anti-GFP antibody. Total protein extracts were immunoblotted. Red asterisks indicate expected band sizes. Numbers on the right indicate kDa values.

To experimentally validate the predicted binding interface between PiE354 and TBC1D15L, we designed two plant expression constructs encoding N- and C-terminal fragments of TBC1D15L. The first construct included the N-terminal RBD fragment, denoted RBDF (1-186), and the second comprised the C-terminal TBC domain fragment, named TBCDF (186-676). Interestingly, the TBCDF construct triggered a slight cell death response, typically noticeable within 3-4 days of transient expression (Figure S2C). Nevertheless, western blot analysis confirmed the successful *in planta* expression of both TBC1D15L fragments, though the protein levels of TBCDF were slightly lower compared to RBDF and full length TBC1D15L (Figure 2B). Our pulldown assays, conducted with protein extracts from *N. benthamiana*, revealed a strong interaction between PiE354 and the RBDF construct. In contrast, we did not detect any interaction between PiE354 and the TBCDF construct. Notably, this finding is in line with our Y2H results (Table S2) and AF2-M predictions (Figure 2A), confirming that PiE354 specifically targets the N-terminal RBD fragment of TBC1D15L.

The AF2-M analysis with stringent parameters identified seven crucial residues on PiE354 (L4, M6, E8, A15, D21, R44, and S81) that are pivotal for its interaction with TBC1D15L (Figures 2C and S2D). To characterize this interaction further, we adopted an alanine scanning strategy, mutating these essential residues in PiE354. Since one of these residues was already encoding alanine (A15), we created a 6A mutant of PiE354 (PiE354^6A^) by substituting the other 6 residues for alanine. Our co-IP assays confirmed that TBC1D15L interacts with PiE354, but not with the PiE354^6A^ mutant or EV (Figure 2D). This result underscores the critical role of these six residues in PiE354 for its interaction with TBC1D15L. Our confocal microscopy analyses further support this notion, showing colocalization of TBC1D15L with PiE354 in puncta, whereas no puncta colocalization was evident with the PiE354^6A^ mutant, mirroring the behavior of the EV control (Figure S2E).

To further elucidate the effector targeting mechanism, we conducted reciprocal mutation experiments focusing on the AF2-predicted binding interface on the host target, TBC1D15L. The AF2-M analysis had previously identified seven crucial residues on TBC1D15L (L100, A135, R115, Y140, A60, L121, and Q124) vital for binding with PiE354 (Figures 2C and S2D). Since A60 and A135 were already encoding alanine, we engineered a mutant, named TBC1D15L^7M^, featuring substitutions of non- alanine residues to alanine and alanine residues to glycine, resulting in a total of seven mutations. We confirmed the expression of GFP:TBC1D15L^7M^ using confocal microscopy, displaying similar characteristics to the WT GFP:TBCD1D15L protein, including cytoplasmic localization and puncta formation (Figure S2F). Our subsequent co-IP assays demonstrated that the TBC1D15L^7M^ mutant was unable to interact with PiE354 (Figure 2E), highlighting the critical role of these seven residues in TBC1D15L for PiE354 targeting. Confocal microscopy analysis provide further support for this notion, as TBC1D15L^7M^ did not colocalize with any of the effectors PiE354, PiE355, and TIKI in punctate structures (Figure S2G). This comprehensive analysis elucidates the binding mechanism between PiE354 and its host target TBC1D15L with key residues in both proteins pinpointed.

### TBC1D15L negatively regulates plant immunity and immune-related secretion in a GAP- dependent manner

Convergence of a conserved *Phytophthora* effector on TBC1D15L hints at a key regulatory role of this RabGAP in immune-related subcellular trafficking. To assess the impact of TBC1D15L on plant immunity, we conducted infection assays with *P. infestans* upon overexpression or silencing of TBC1D15L. The dual catalytic fingers of TBC1D15L, crucial for stimulating GTP hydrolysis of Rab GTPases, are located at R446 and Q483 positions within the TBC domain (Figure 2A). We created the TBC1D15L GAP mutant (TBC1D15L^GAP^) by the dual mutations at R446A and Q483A positions, which typically impair the GAP activity of TBC-containing RabGAP proteins^23^. AF2 modeling revealed that the WT and GAP mutant of TBC1D15L maintain a high level of structural similarity, indicated by a low RMSD value of 0.286 (Figures 3A and S3A), suggesting that the overall protein architecture of the TBC1D15L^GAP^ mutant is not perturbed. Both GFP:TBC1D15L and GFPTBC1D15L^GAP^ successfully pulled down RFP:TIKI from *N. benthamiana* protein extracts, but not the RFP:EV control (Figure S3B). Reciprocal co-IP experiments using RFP fusions of TBC1D15L constructs with GFP:TIKI further validated the specific interaction between the effector and the RabGAP protein (Figure S3B). These results provide strong evidence that TIKI associates with TBC1D15L *in planta* independent of the GAP activity of TBC1D15L.

**Figure 3.**
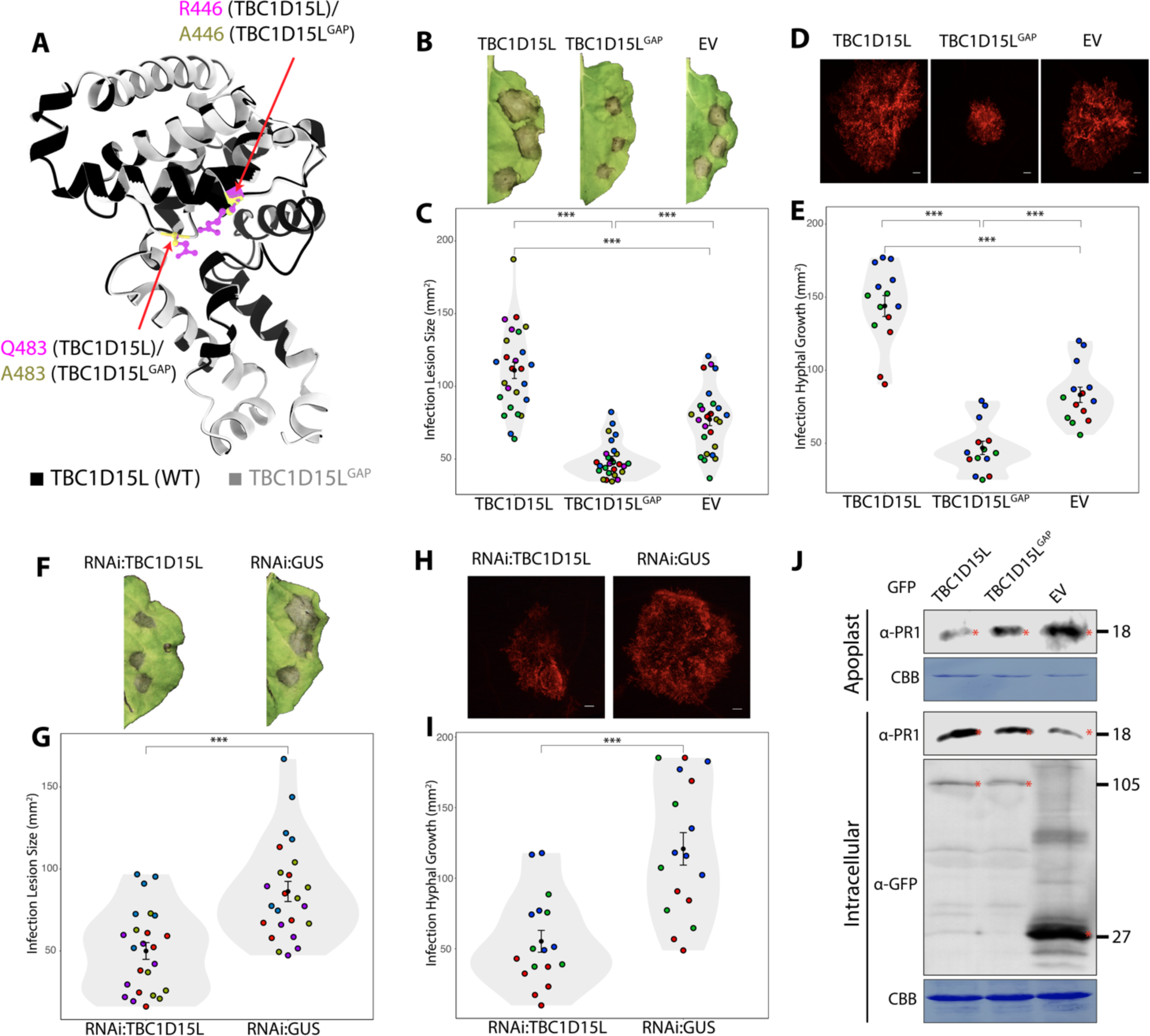
TBC1D15L negatively regulates plant immunity through its GAP function. (A) Structural alignment of TBC1D15L and its GAP mutant TBC1D15L^GAP^. Structural predictions were obtained via AF2. The model shows conservation of the overall protein structure, with RMSD value being 0.286. (B-C) TBC1D15L increases susceptibility to *P. infestans* in a GAP activity dependent manner. (B) *N. benthamiana* leaves expressing TBC1D15L, TBC1D15L^GAP^ or EV control were infected with *P. infestans*, and pathogen growth was calculated by measuring infection lesion size at 8 days post- inoculation. (C) TBC1D15L expression (111.0 mm^2^, N = 78) significantly increases *P. infestans* lesion size compared to EV control (77.3 mm^2^, N = 78), while TBC1D15L^GAP^ expression (49.5 mm^2^, N = 78) significantly reduces the lesion size compared to EV control. (D-E) TBC1D15L increases *P. infestans* hyphal growth in a GAP activity dependent manner. (D) *N. benthamiana* leaves expressing TBC1D15L, TBC1D15L^GAP^ or EV control were infected with tdTomato-expressing *P. infestans*, and pathogen growth was calculated by measuring hyphal growth using fluorescence stereomicroscope at 5 days post-inoculation. Scale bars, 10 μm. (E) TBC1D15L expression (144.0 mm^2^, N = 42) significantly increases *P. infestans* hyphal growth compared to EV control (83.2 mm^2^, N = 42), while TBC1D15L^GAP^ expression (46.9 mm^2^, N = 42) significantly reduces the hyphal growth compared to EV control. (F-G) Silencing *TBC1D15L* reduces susceptibility to *P. infestans*. (F) *N. benthamiana* leaves expressing RNAi:TBC1D15L, or RNAi:GUS control were infected with *P. infestans*, and pathogen growth was calculated by measuring infection lesion size at 8 days post-inoculation. (G) RNAi:TBC1D15L expression (50.0 mm^2^, N = 72) significantly reduces *P. infestans* lesion size compared to RNAi:GUS control (86.3 mm^2^, N = 72). (H-I) Silencing *TBC1D15L* reduces hyphal growth of *P. infestans*. (H) *N. benthamiana* leaves expressing RNAi:TBC1D15L, or RNAi:GUS control were infected with tdTomato-expressing *P. infestans*, and pathogen growth was calculated by measuring hyphal growth using fluorescence stereomicroscope at 5 days post-inoculation. Scale bars, 10 μm. (I) RNAi:TBC1D15L expression (55.3 mm^2^, N = 51) significantly reduces *P. infestans* lesion size compared to RNAi:GUS control (120.8 mm^2^, N = 51). Each color represents an independent biological replicate. Each dot represents the average of 3 infection spots on the same leaf. Statistical differences were analyzed by Student’s t-test or Mann-Whitney U test in R. Measurements were highly significant when p<0.001 (***). (J) *N. benthamiana* leaves were agroinfiltrated to express GFP:TBC1D15L, GFP:TBC1D15L^GAP^, or GFP:EV. The infiltrated leaves were challenged with *P. infestans* extract at 3 dpi and proteins were extracted from the apoplast and leaf tissue at 4 dpi and immunoblotted. Western blot shows TBC1D15L, but not its GAP mutant TBC1D15L^GAP^, reduces antimicrobial PR1 secretion into the apoplast, compared to EV control. Red asterisks show expected band sizes. Numbers on the right indicate kDa values.

Across five independent experiments, we observed that overexpression of TBC1D15L consistently enhanced *P. infestans* infection symptoms, notably increasing lesion size, compared to the EV control (Figures 3B and 3C). Intriguingly, overexpression of the GAP mutant TBC1D15L^GAP^ led to a significant reduction in infection lesion size relative to the EV control (Figures 3B and 3C), suggesting a functional link between the GAP activity of TBC1D15L and its role in plant immunity. To further substantiate the adverse effect of TBC1D15L on immunity, we performed additional infection assays using the red fluorescent *P. infestans* strain 88069td. This approach allows for the direct quantification of pathogen biomass by measuring biotrophic hyphal growth using fluorescence microscopy. Parallel to our earlier infection assays (Figures 3B and 3C), elevating TBC1D15L levels significantly boosted *P. infestans* hyphal growth in three independent experiments (Figures 3D and 3E). Conversely, the overexpression of the GAP mutant TBC1D15L^GAP^ led to a marked decrease in pathogen hyphal growth compared to the EV control (Figures 3D and 3E), indicating that the GAP activity of TBC1D15L is crucial for facilitating disease susceptibility.

To complement the overexpression assays, we performed RNA interference (RNAi) to downregulate *TBC1D15L* gene expression in *N. benthamiana* using a hairpin RNAi construct (RNAi:TBC1D15L) (Figure S3C). Consistent with our overexpression assays, which hinted at a negative role for TBC1D15L in plant immunity, silencing of TBC1D15L significantly reduced *P. infestans* infection lesions compared to the silencing control, RNAi:GUS (Figures 3F and 3G). In addition, silencing of TBC1D15L also reduced *P. infestans* hyphal growth compared to the GUS silencing control (Figures 3H and 3I). Collectively, these results further confirm the role of TBC1D15L as a negative regulator of plant immunity. This notion is supported by the observed dominant negative phenotype of the TBC1D15L^GAP^ mutant, which enhances resistance.

Given the regulatory role of RabGAPs in subcellular trafficking and our finding that TBC1D15L negatively regulates plant immunity via its GAP function, we reasoned that TBC1D15L might be controlling immune-related secretion. To test this hypothesis, we analyzed the potential impact of TBC1D15L on defense-related secretion by monitoring the native levels of pathogenesis-related protein 1 (PR1) in the apoplast. We measured PR1 levels by using a specific antibody raised against it following overexpression of TBC1D15L, TBC1D15L^GAP^ mutant, or the GFP:EV control. We noted a drastic decrease in PR1 secretion into the apoplast when TBC1D15L was overexpressed, in contrast to the TBC1D15L^GAP^ mutant or EV control overexpression (Figure 3J). This outcome strongly supports the notion that TBC1D15L subverts plant immunity by suppressing defense-related secretion.

### PiE354 co-opts TBC1D15L to subvert plant immunity and defense-related secretion

To elucidate the functional relationship between PiE354 and TBC1D15L, we first determined the extent to which PiE354 or its mutant PiE354^6A^—impaired in binding to its host target TBC1D15L (Figure 2D)— influence susceptibility to *P. infestans.* We measured this through infection assays on *N. benthamiana* leaf patches that transiently express PiE354, PiE354^6A^, or EV control. In four independent infection assays, we observed a consistent increase in *P. infestans* infection lesion size in leaf patches expressing PiE354 compared to EV control (Figures 4A and 4B). In contrast, leaf samples expressing PiE354^6A^ mutant did not exhibit any increase in infection lesion size relative to the EV control (Figures 4A and 4B). These findings demonstrate that PiE354 enhances *P. infestans* virulence on plants and its ability to exacerbate *P. infestans* virulence is linked to its interaction with TBC1D15L.

**Figure 4.**
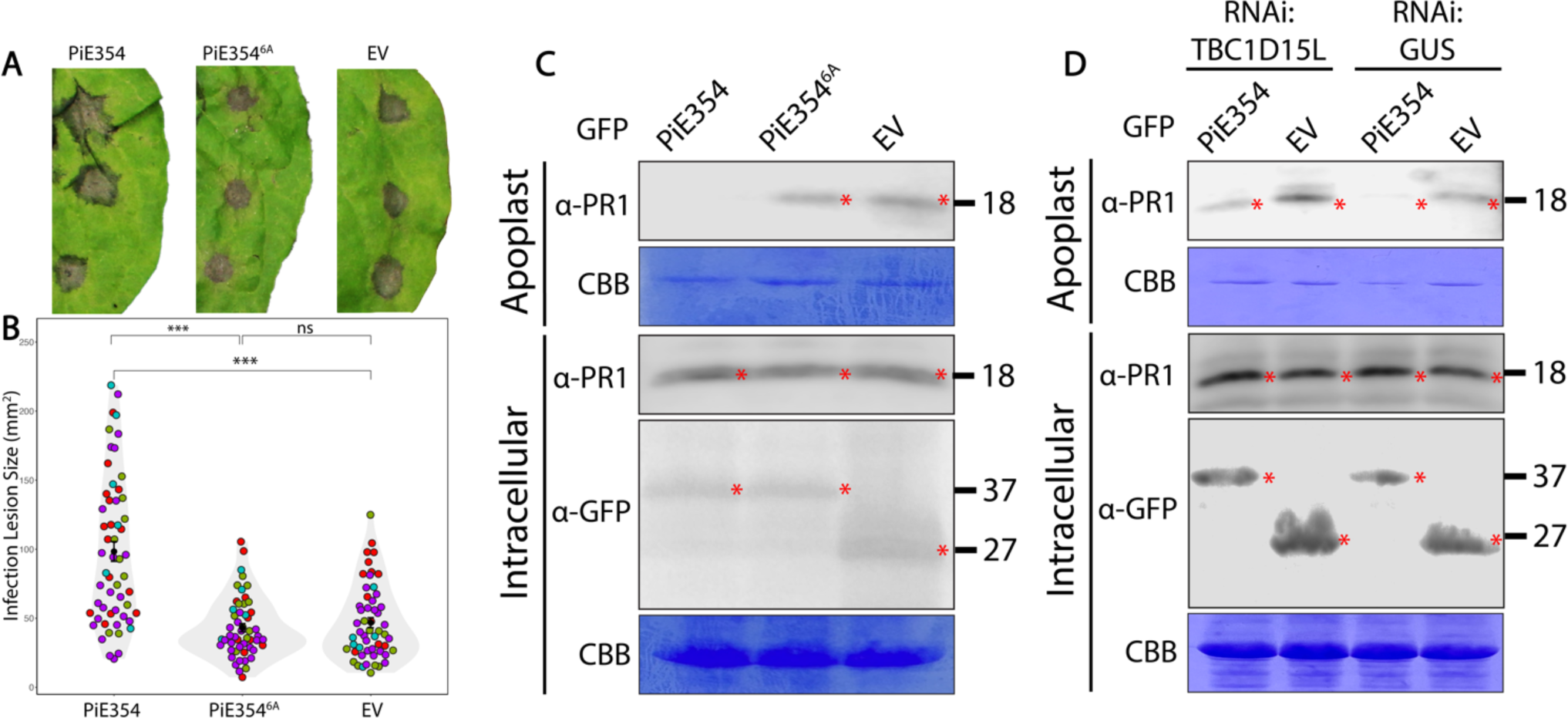
PiE354 co-opts TBC1D15L to subvert plant immunity and defense-related secretion. (A-B) PiE354 enhances susceptibility to *P. infestans*, contingent on its interaction with TBC1D15L. (A) *N. benthamiana* leaves expressing PiE354, PiE354^6A^, or EV control were infected with WT *P. infestans*, and pathogen growth was calculated by measuring infection lesion size at 8 days post-inoculation. (B) PiE354 expression (98.3 mm^2^, N = 56) significantly increases *P. infestans* lesion size compared to EV control (43.2 mm^2^, N = 52), while the expression of PiE354^6A^ (46.8 mm^2^, N = 54) has no significant effect compared to EV control. Each color represents an independent biological replicate. Each dot represents an infection spot. Statistical differences were analyzed by Mann- Whitney U test in R. Measurements were highly significant when p<0.001 (***). (C-D) For PR1 secretion assays, the infiltrated leaves were challenged with *P. infestans* extract at 3 dpi and proteins were extracted from the apoplast and leaf tissue at 4 dpi and immunoblotted. (C) Western blot shows PiE354 suppresses antimicrobial PR1 secretion into the apoplast, dependent on the interaction with its host target TBC1D15L. *N. benthamiana* leaves were infiltrated to express GFP:PiE354, GFP:PiE354^6A^ (mutant that lacks the capability to interact with TBC1D15L), or GFP:EV. (D) Western blot shows the ability of PiE354 to inhibit PR1 secretion is less pronounced when TBC1D15L is silenced, compared to GUS-silenced control condition. *N. benthamiana* leaves were infiltrated to express RNAi:TBC1D15L or RNAi:GUS control, with either PiE354 or EV control. Red asterisks show expected band sizes. Numbers on the right indicate kDa values.

We then investigated whether PiE354 similarly affects PR1 secretion in *N. benthamiana,* similar to the effects observed with TBC1D15L overexpression (Figure 3J). Heterologous expression of PiE354, but not the EV control, effectively suppressed PR1 secretion into the apoplast (Figures 4C and S4). In contrast, PiE354^6A^ mutant did not have any effect on PR1 secretion, behaving much like the EV control (Figure 4C). This suggests that PiE354 mirrors the effects of TBC1D15L overexpression, reinforcing the notion that it facilitates the GAP function of TBC1D15L to hinder defense-related secretion (Figure 3J).

Lastly, to ascertain the role of PiE354 in disrupting defense-related secretion through its interaction with TBC1D15L, we conducted PR1 secretion assays in *TBC1D15L*-silenced plants. As expected, in leaf patches with RNAi:GUS silencing control, PiE354 effectively reduced PR1 secretion into the apoplast, relative to the EV (Figure 4D). Conversely, when *TBC1D15L* was silenced, the capacity of PiE354 to inhibit PR1 secretion was less pronounced, although there was still a noticeable reduction in PR1 secretion compared to the EV (Figure 4D). This is a reasonable outcome given the RNAi:TBC1D15L construct does not fully deplete the TBC1D15L transcripts (Figure S3C). These findings demonstrate that the capacity of PiE354 to interfere with defense-related secretion and plant immunity is intricately linked to TBC1D15L.

### Rab8a, a Rab GTPase that mediates defense-related secretion, is a GAP substrate of TBC1D15L

We next focused on determining the cognate Rab GTPase partner of TBC1D15L. Through an IP-MS interactome screen, we identified Rab8a as a candidate Rab substrate of TBC1D15L (Table S4). Given the previous findings by us and others that Rab8a plays a positive role in plant immunity against *P. infestans* and mediates PR1 secretion^3,14,24^, we reasoned that Rab8a could be the cognate partner of TBC1D15L.

To corroborate our IP-MS findings, we set out to verify the interaction between Rab8a and TBC1D15L through co-IPs. These assays confirmed that RFP:TBC1D15L interacts with GFP:Rab8a but not with GFP:EV. Also, RFP:EV control did not interact with GFP:Rab8a, indicating that TBC1D15L specifically binds GFP:Rab8a (Figure 5A). Our confocal microscopy analysis also showed that TBC1D15L, but not the EV control, colocalizes with Rab8a in puncta (Figure 5B).

**Figure 5.**
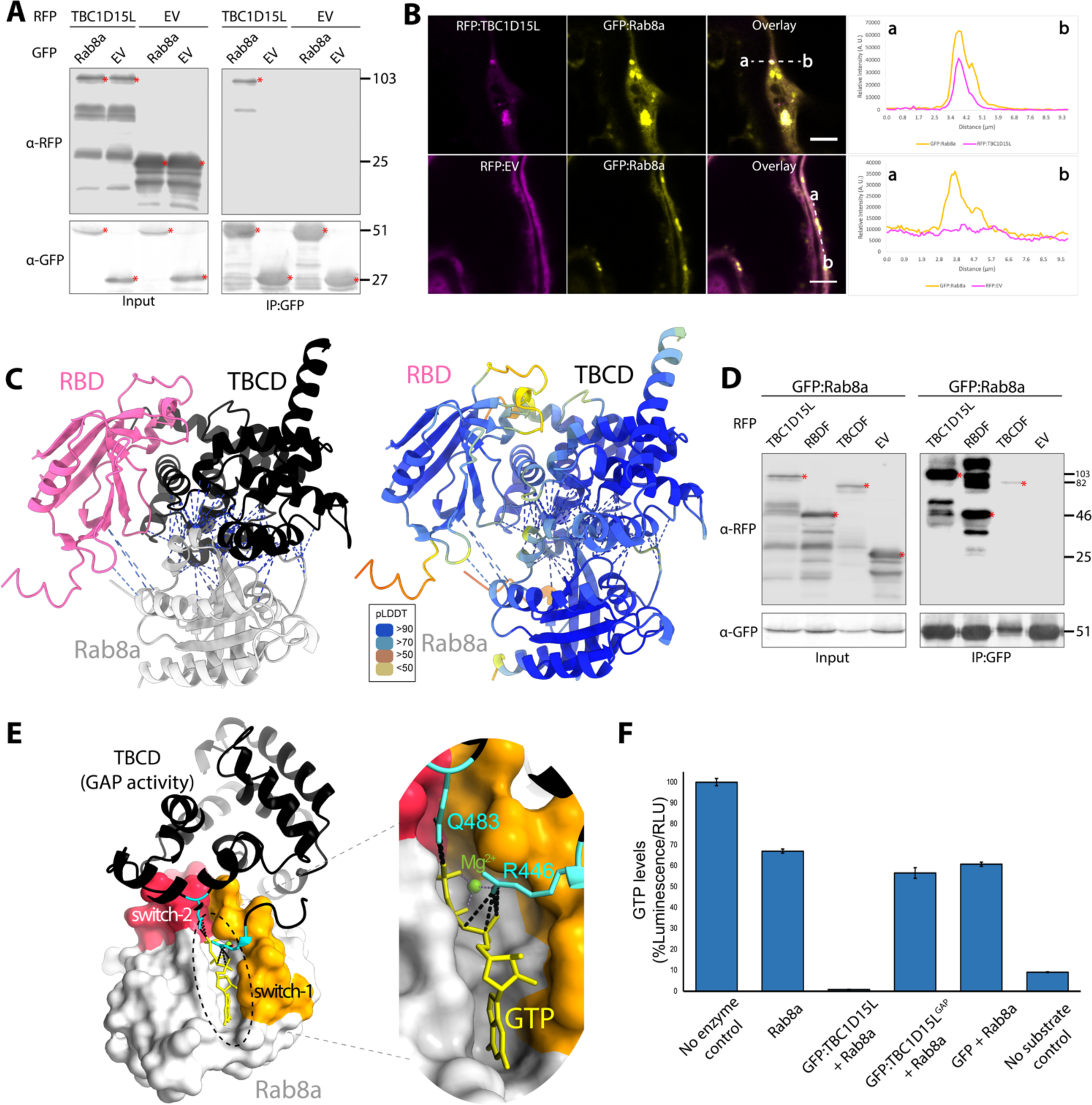
Rab8a is a GAP substrate of TBC1D15L. (A) TBC1D15L, but not the EV control, interacts with Rab8a *in planta*. RFP:TBC1D15L was transiently co-expressed with either GFP:Rab8a, or the GFP:EV control. RFP:EV was used as a control for RFP:TBC1D15L. IPs were obtained with anti-GFP antibody. Total protein extracts were immunoblotted. Red asterisks indicate expected band sizes. Numbers on the right indicate kDa values. (B) Rab8a colocalizes with TBC1D15L in puncta *in planta*. Confocal micrographs of *N. benthamiana* leaf epidermal cells transiently expressing either RFP:TBC1D15L (1^st^ row), or RFP:EV (2^nd^ row), with GFP:Rab8a. Presented images are single plane images. Transects in overlay panels correspond to line intensity plots depicting the relative fluorescence across the marked distance. Scale bars, 5 µm. (C) AF2-M modeling of Rab8a with individual RBD and TBCD of TBC1D15L in complex. (Left panel) Rab8a interacts with both the RBD and TBCD of TBC1D15L. (Right panel) The colors of the AF2-M model are based on the AF2-calculated prediction confidence score (pLDDT) as indicated in the rectangular box. (D) Rab8a interacts with the full length, N-terminal RBD fragment (RBDF) and C-terminal TBCD fragment (TBCDF) of TBC1D15L, but not with the EV control. GFP:Rab8a was transiently co-expressed with either RFP:TBC1D15L, RFP:RBDF, RFP:TBCDF, or RFP:EV. IPs were obtained with anti-GFP antibody. Total protein extracts were immunoblotted. Red asterisks indicate expected band sizes. Numbers on the right indicate kDa values. (E) Predicted AF2-M model of Rab8a in complex with full length TBC1D15L, focusing on TBCDF of TBC1D15L. The catalytic dual fingers of the TBCD, R446 and Q483 (colored cyan), are favorably positioned across the GTP binding pocket of Rab8a, flanked by switch-1 (pink) and switch-2 (orange) regions, and making contacts with the GTP molecule within the pocket. (F) TBC1D15L stimulates the GTPase activity of Rab8a, dependent on its GAP activity. A luciferase-based GTPase assay (GTPase-Glo™ Assay kit by Promega) was employed to quantify the amount of GTP levels over a 120-minute duration at room temperature. The bar graph illustrates the effect of TBC1D15L, TBC1D15L^GAP^, or GFP control on the GTPase activity of Rab8a across three technical repeats. No enzyme control does not contain Rab8a. No substrate control does not contain GTP.

The AF2-M prediction of the TBC1D15L-Rab8a complex revealed interactions between Rab8a and both the RBD and TBC domains of TBC1D15L, with the binding interface predominantly oriented towards the TBC domain (Figure S5A). This bias could be due to the presence of available GAP-Rab crystal structures in the mammalian field and the higher sequence conservation of the TBC domain compared to the RBD, resulting in higher confidence score models for the TBC domain binding interface than the RBD interface. AF2-M modeling of Rab8a with individual RBD and TBC domain of TBC1D15L showed high-confidence interactions between each domain of TBC1D15L and Rab8a (Figure 5C). To determine if Rab8a engages both the RBD and TBC domain as suggested by the AF2- M models, we performed pulldown assays using Rab8a with the two TBC1D15L fragments. Remarkably, our co-IP results revealed that Rab8a interacts with both the RBD and TBCD fragments of TBC1D15L (Figure 5D), corroborating the AF2-M model predictions. However, it is difficult to discern whether Rab8a binds both domains simultaneously or alternates between them. To further characterize the binding mechanism between Rab8a and TBC1D15L, we investigated whether their interaction is dependent on the GAP activity of TBC1D15L. Co-IP and western blot analysis showed that Rab8a interacts with both WT TBC1D15L and its GAP mutant, but not with the EV control, indicating that Rab8a interacts with TBC1D15L independently of the GAP function of TBC1D15L (Figure S5B). Our confocal microscopy analysis also revealed that both TBC1D15L and its GAP mutant, but not the EV control, colocalize with Rab8a (Figures 5B and S5C). These findings indicate that while the RBD and TBCD of TBC1D15L facilitate its association with Rab8a, the catalytic function of the GAP domain is dispensable for Rab8a binding.

Having validated the TBC1D15L-Rab8a interaction, we next investigated the functional interplay between the two proteins, focusing on investigating whether Rab8a is a GAP substrate of TBC1D15L. We first employed AF2-M to visualize the interaction between TBC1D15L and Rab8a, with a specific focus on the TBCD fragment that contains the GAP activity. AF2-M prediction of the Rab8a-TBC1D15L complex revealed a high-confidence model in which the TBCD makes multiple contacts with the switch-1 and switch-2 regions of Rab8a, which are flanking the GTP binding pocket of Rab8a and is crucial for regulating its GTP hydrolysis activity (Figure S5D). We then introduced GTP inside the Rab8a GTP pocket on the AF2-M model by using the crystal structure of human Rab8a bound to GTP (PDB: 6WHE) as described before^24^.The resulting model revealed that the catalytic dual fingers of the TBC domain, specifically R446 and Q483, are favorably positioned to establish contacts with the GTP molecule within the Rab8a GTP-binding pocket (Figure 5E). This high-confidence AF2-M model suggests a reasonable configuration of the GAP domain to catalyze the conversion of Rab8a-GTP to Rab8a-GDP.

To experimentally determine whether Rab8a is a substrate of TBC1D15L, we conducted on beads GAP activity assays. We isolated total protein extracts from *N. benthamiana* leaves expressing GFP:TBC1D15L, GFP:TBC1D15L^GAP^ mutant, or GFP:EV control and concentrated them on GFP-trap beads via immunoprecipitation. Prior to assessing the activities of the immobilized constructs on beads, we confirmed the functionality of the GTPase activity of Rab8a purified from *E. coli* (Figure S5E). Next, we incubated purified Rab8a alone, or with GFP:TBC1D15L, GFP:TBC1D15L^GAP^, or the GFP:EV control. Rab8a alone induced a 30-35% reduction in GTP levels compared to the buffer with no enzyme control within two hours. Remarkably, incubation of Rab8a with the affinity resin that pulled down GFP:TBC1D15L completely depleted GTP levels (Figure 5F). In contrast, incubation with GFP:TBC1D15L^GAP^ or the EV control did not considerably alter GTP levels compared to Rab8a alone (Figure 5F). These findings conclusively show that TBC1D15L substantially enhances the GTP hydrolysis activity of Rab8a, acting as a canonical GAP, and this activity is reliant on the functional integrity of its TBC domain.

### TBC1D15L negatively regulates immunity by restricting Rab8a-mediated subcellular trafficking towards the cell surface

Having determined the *in vitro* GAP activity of TBCD15L on Rab8a, we next sought to determine how TBC1D15L regulates Rab8a functions *in vivo*. Our previous work showed that Rab8a localizes with a more pronounced signal at the plasma membrane compared to the vacuolar membrane (tonoplast), indicating a predominant Rab8a transport route towards the cell surface^14^. To investigate if TBC1D15L modulates subcellular localization of Rab8a, we used confocal microscopy. Overexpression of RFP:TBC1D15L resulted in a marked reduction of the plasma membrane levels of GFP:Rab8a with respect to the tonoplast, indicating the redistribution of GFP:Rab8a trafficking towards the vacuole (Figures 6A and 6B). In contrast, co-expression with either TBC1D15L^GAP^ mutant or the EV control maintained the primary localization of GFP:Rab8a at the plasma membrane (Figures 6A and 6B), aligning with our previous findings^14^. These results provide compelling evidence that TBC1D15L regulates Rab8a-mediated trafficking from the cell surface to the vacuole, further affirming its role as a GAP for Rab8a. Consistent with prior research showing the role of Rab8a in PR1 release into the apoplast^3^, we infer that TBC1D15L hinders PR1 secretion (Figure 3J) by redirecting Rab8a-mediated trafficking away from the cell surface.

**Figure 6.**
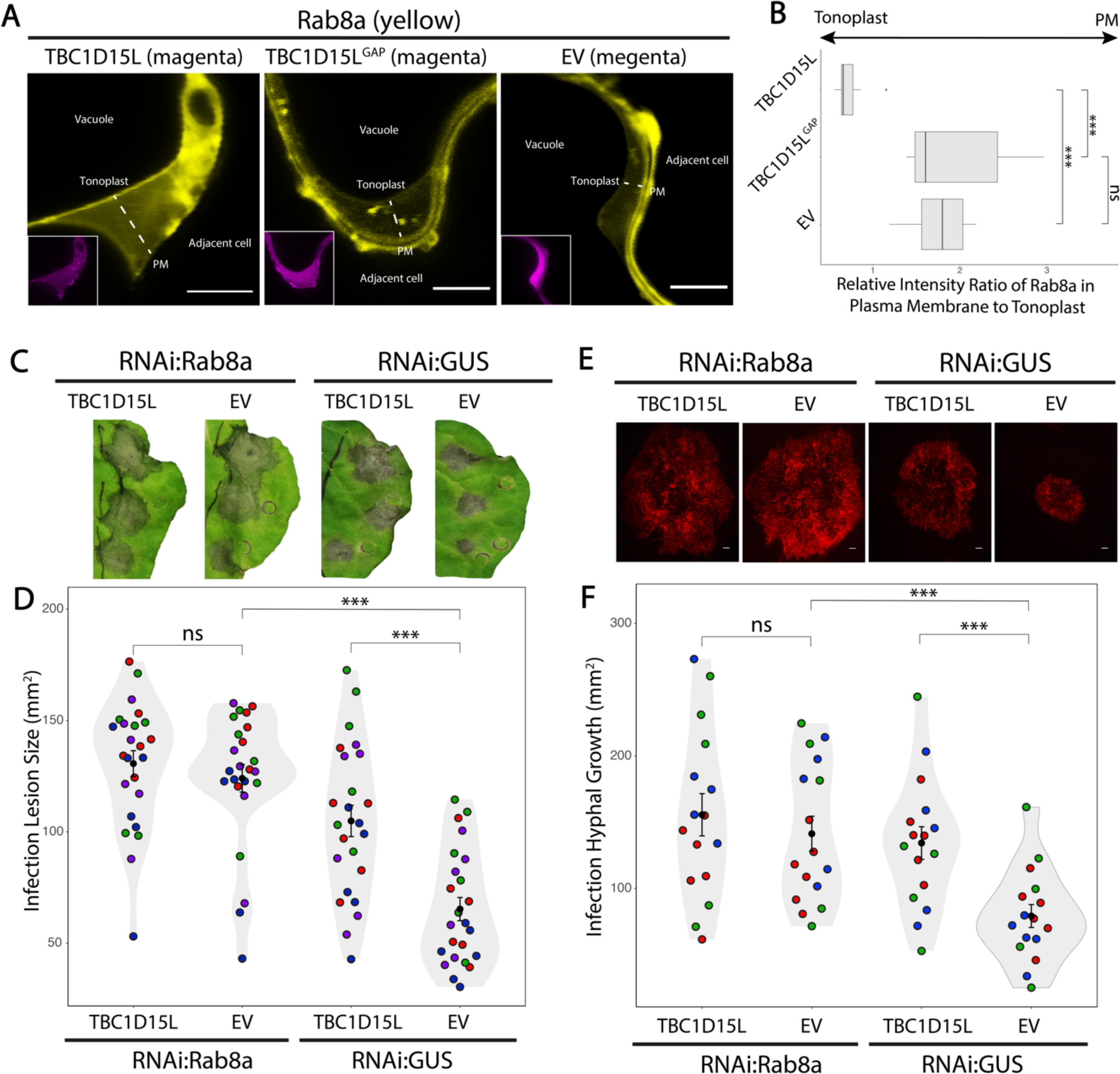
TBC1D15L negatively regulates immunity by restricting Rab8a-mediated subcellular trafficking towards the cell surface. (A-B) TBC1D15L diverts Rab8a localization from the plasma membrane to the tonoplast dependent on its GAP activity. (A) Confocal micrographs of *N. benthamiana* leaf epidermal cells transiently co-expressing either RFP:TBC1D15L, RFP:TBC1D15L^GAP^, or RFP:EV, with GFP:Rab8a. Presented images are single plane images. Scale bars, 5 µm. (B) Box plot illustrates that TBC1D15L expression (0.74, N = 8) significantly reduces the relative intensity ratio of Rab8a in plasma membrane to tonoplast compared to the EV control (1.76, N = 8), while TBC1D15L^GAP^ expression (1.94, N = 8) has no significant effect compared to the EV control. (C-D) The significant increase in *P. infestans* lesion size caused by TBC1D15L expression is negated when Rab8a is silenced. (C) RNAi:Rab8a, or RNAi:GUS control was co-expressed with either TBC1D15L, or EV control in WT leaves. The agroinfiltrated leaves were infected with WT *P. infestans*, and pathogen growth was calculated by measuring infection lesion size at 7 days post-inoculation. (D) In RNAi:Rab8a leaves, TBC1D15L expression (130.6 mm^2^, N = 72) has no significant effect on *P. infestans* lesion size compared to the EV control (124.0 mm^2^, N = 72). In RNAi:GUS control leaves, TBC1D15L expression (104.9 mm^2^, N = 72) significantly increases *P. infestans* lesion size compared to the EV control (65.3 mm^2^, N = 72). (E-F) The significant increase in *P. infestans* hyphal growth caused by TBC1D15L expression is negated when Rab8a is silenced. (E) RNAi:Rab8a, or RNAi:GUS control was co-expressed with either TBC1D15L, or EV control in WT leaves. The agroinfiltrated leaves were infected with tdTomato-expressing *P. infestans,* and pathogen growth was calculated by measuring hyphal growth using fluorescence stereomicroscope at 5 days post-inoculation. Scale bars, 10 µm. (F) In RNAi:Rab8a leaves, TBC1D15L expression (155.6 mm^2^, N = 48) has no significant effect on *P. infestans* hyphal growth compared to the EV control (141.3 mm^2^, N = 48). In RNAi:GUS control leaves, TBC1D15L expression (134.2 mm^2^, N = 48) significantly increases *P. infestans* hyphal growth compared to the EV control (79.1 mm^2^, N = 48). Each color represents an independent biological replicate. Each dot represents the average of 3 infection spots on the same leaf. All statistical differences were analyzed by Student’s T-test, or Mann-Whitney U test in R. Measurements were highly significant when p<0.001 (***).

Considering previous studies demonstrating the role of Rab8a in plant immunity, we hypothesized that the negative impact of TBC1D15L on immunity might be due to its influence on Rab8a trafficking. To elucidate the interplay between TBC1D15L and Rab8a in plant immunity, we silenced Rab8a using a hairpin RNAi construct^14^, and examined whether TBC1D15L can still suppress immunity. Consistent with the well-established defense roles of Rab8a against *P. infestans*^3,14^, silencing Rab8a increased infection lesion sizes compared to the GUS-silencing control. In GUS-silenced plants, TBC1D15L overexpression significantly enlarged infection lesions relative to the EV control, aligning with our findings of TBC1D15L negatively impacts immunity (Figures 6C and 6D). Conversely, in Rab8a- silenced plants, TBC1D15L overexpression did not significantly alter infection lesion sizes compared to the EV control (Figures 6C and 6D). This pattern was also evident when measuring pathogen biomass in infection assays using the red fluorescent *P. infestans* strain 88069td (Figures 6E and 6F). These results collectively indicate that the negative influence of TBC1D15L on plant immunity is dependent on its regulation on Rab8a.

### PiE354 diverts Rab8a trafficking by hijacking the TBC1D15L-Rab8a complex

AF2-M modeling of the PiE354-Rab8a-TBC1D15L complex reveals a compelling tripartite interaction (Figures 7A and S6), aligning with our protein-protein interaction assays (Figures 2B and 5D). It appears that PiE354 alters the orientation of Rab8a towards the TBCD fragment harboring the GAP activity while associating with the RBD interface (Figure 7A). This is supported by the loss of AF2-M- predicted contacts between RBD and Rab8a (Figure 5C) when PiE354 is present (Figure 7A).

**Figure 7.**
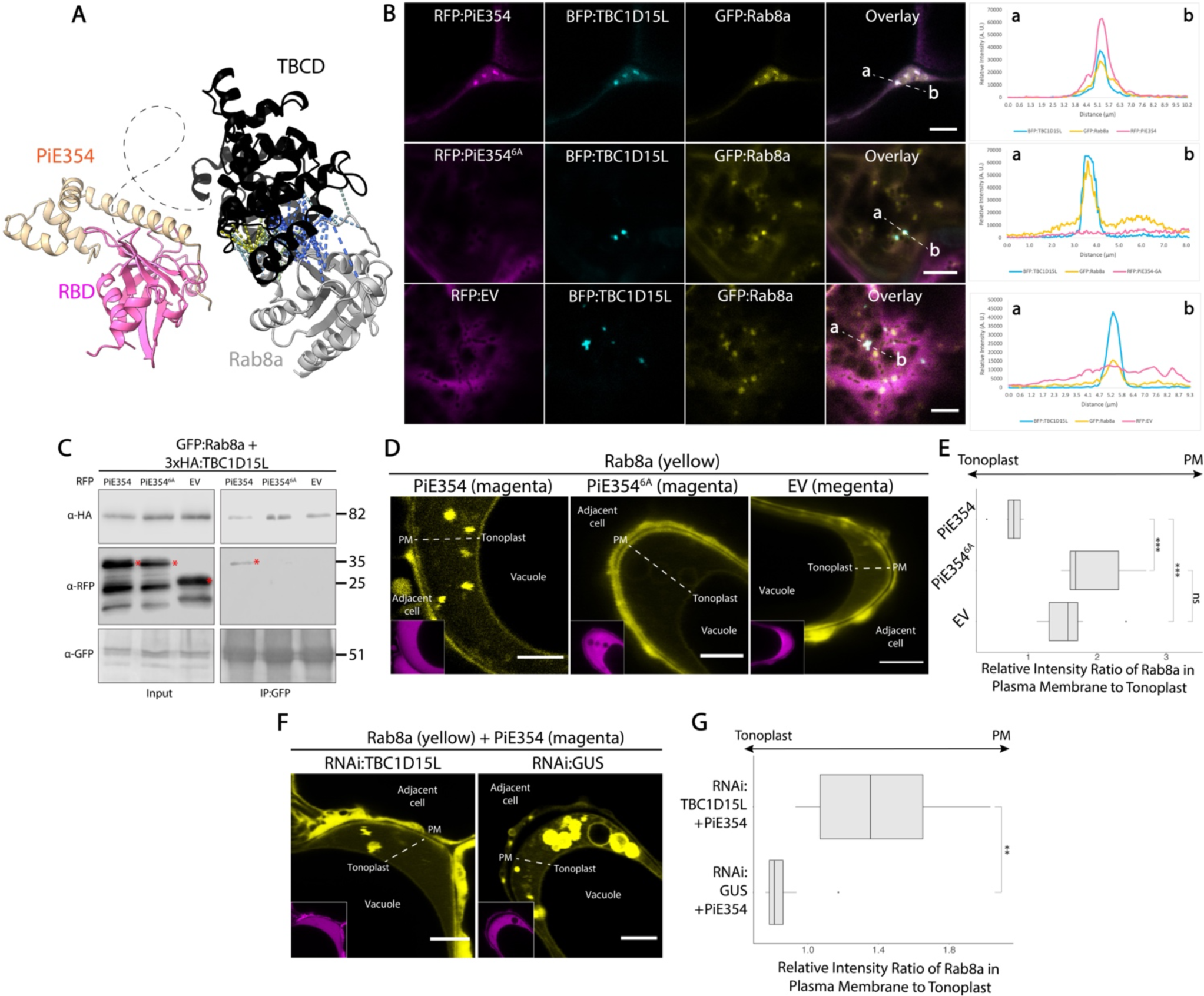
PiE354 targets the TBC1D15L-Rab8a complex and re-routes Rab8a trafficking towards the vacuole. (A) AF2-M-predicted model of PiE354 in complex with the TBC1D15L-Rab8a pair. The model reveals that in the presence of PiE354, Rab8a and the RBD of TBC1D15L are no longer interacting, as PiE354 tilts Rab8a towards the functional TBCD of TBC1D15L. (B) PiE354, but not PiE354^6A^ or EV, colocalizes with Rab8a and TBC1D15L altogether in puncta. Confocal micrographs of *N. benthamiana* leaf epidermal cells transiently co-expressing either RFP:PiE354 (1^st^ row), RFP:PiE54^6A^ (2^nd^ row), or RFP:EV (3^rd^ row), with BFP:TBC1D15L and GFP:Rab8a. Presented images are single plane images. Transects in overlay panels correspond to line intensity plots depicting the relative fluorescence across the marked distance. Scale bars, 5 µm. (C) PiE354 binds to the TBC1D15L-Rab8a pair dependent on its interaction with TBC1D15L. GFP:Rab8a and 3xHA:TBC1D15L were transiently co-expressed with either RFP:PiE354, RFP:PiE354^6A^, or RFP:EV. IPs were obtained with anti-GFP antibody. Total protein extracts were immunoblotted. Red asterisks indicate expected band sizes. Numbers on the right indicate kDa values. (D-E) PiE354 diverts Rab8a localization from the plasma membrane to the tonoplast dependent on its interaction with TBC1D15L. (D) Confocal micrographs of *N. benthamiana* leaf epidermal cells transiently expressing either RFP:PiE354, RFP:PiE354^6A^, or RFP:EV, with GFP:Rab8a. Presented images are single plane images. Scale bars, 5 µm. (E) Box plot showing the expression of PiE354 (0.77, N = 8), but not PiE354^6A^ which lacks binding capability to TBC1D15L (1.93, N = 8), significantly reduces the relative intensity ratio of Rab8a in plasma membrane to tonoplast compared to EV control (1.59, N = 8). EV and PiE354^6A^ result in a predominant localization of Rab8a to the plasma membrane, while PiE354 re-directs the predominant localization of Rab8a to the tonoplast. (F-G) Silencing *TBC1D15L* nullifies the ability of PiE354 diverting Rab8a localization from plasma membrane to the tonoplast. (F) Confocal micrographs of *N. benthamiana* leaf epidermal cells transiently expressing either RNAi:TBC1D15L, or RNAi:GUS control, with RFP:PiE354 and GFP:Rab8a. Presented images are single plane images. Scale bars, 5 µm. (G) Box plot illustrating under *TBC1D15L*-silenced condition, PiE354 expression (1.40, N = 8) leads to a significant increase in the relative intensity ratio of Rab8a in plasma membrane to tonoplast, compared to the GUS-silenced control condition (0.86, N = 8). All statistical differences were analyzed by Mann-Whitney U test in R. Measurements were significant when p<0.01 (**), and highly significant when p<0.001 (***).

To explore the potential tripartite interaction of PiE354, TBC1D15L and Rab8a, we first performed colocalization assays using confocal microscopy. In agreement with the AF2-M prediction of the complex mediated by TBCD15L, TBC1D15L and Rab8a colocalized in punctate structures with PiE354, but not with the PiE354^6A^ mutant or the EV control (Figure 7B). To gain further evidence that PiE354 can form a complex with TBC1D15L and Rab8a, we next performed co-IPs using protein extracts from *N. benthamiana* leaves expressing GFP:Rab8a and 3xHA:TBC1D15L with either RFP:PiE354, RFP:PiE354^6A^ mutant or the RFP:EV control. GFP pull-down assays showed that GFP:Rab8a pulls-down both 3xHA:TBC1D15L and RFP:PiE354, but not the RFP:PiE354^6A^ mutant or RFP:EV (Figure 7C). These results are in agreement with the predicted AF2-M models (Figure 7A) and outputs of the colocalization assays (Figure 7B), reinforcing the view that PiE354 targets the TBC1D15L-Rab8a complex. Since Rab8a did not pull-down the PiE354^6A^ mutant, which cannot bind TBC1D15L, we conclude that Rab8a-PiE354 interaction is mediated by TBC1D15L.

Building on these insights, we hypothesized that PiE354 targets TBC1D15L to leverage its GAP activity for disrupting Rab8a-mediated trafficking. To determine if PiE354 mirrors the effects of TBC1D15L overexpression, specifically in redirecting Rab8a trafficking towards the vacuole, we conducted detailed observations using confocal microscopy. Quantitative image analysis showed that when co-expressed with PiE354, Rab8a predominantly localized to the tonoplast, diverging from its usual plasma membrane localization^14^ observed with the PiE354^6A^ mutant or the EV control (Figures 7D and 7E). This finding aligns with our hypothesis, suggesting that PiE354 mimics the effect of TBC1D15L overexpression, promoting the deactivation of Rab8a by TBC1D15L. This notion is further reinforced by our experiments on the diversion of Rab8a trafficking by PiE354 following the silencing of *TBC1D15L*. Quantitative analysis of confocal micrographs from these experiments revealed that in *TBC1D15L*-silenced leaf patches, but not in GUS-silenced control leaves, PiE354 was unable to redirect Rab8a toward the tonoplast (Figures 7F and 7G). These results, combined with the fact that the PiE354^6A^ mutant, which cannot bind TBC1D15L, does not alter the trafficking of Rab8a, suggest that PiE354 potentiates the GAP activity of TBC1D15L, ultimately diverting Rab8a trafficking from the cell surface to the vacuole. These findings are in line with our earlier results showing diminished PR1 secretion caused by the PiE354-TBC1D15L complex (Figures 4C and 4D), showing that PiE354 engages TBC1D15L to divert Rab8a-mediated trafficking critical for defense at the pathogen interface.

### A plant pathogen effector that redirects defense-related secretion by co-opting a key transport regulator

Our results suggest a model where PiE354, along with its homologs PiE355 and TIKI, harness the GAP function of TBC1D15L on Rab8a, a key Rab GTPase involved in antimicrobial secretion. This process reroutes Rab8a-mediated antimicrobial secretion away from the pathogen interface and towards the vacuole, thereby hindering the ability of the plant to mount an effective immune response (Figure 8A). We propose a mechanistic model in which PiE354 binds to the RBD of TBC1D15L, propelling Rab8a towards the TBCD of TBC1D15L, harboring the GAP function. This triggers an accelerated GTP hydrolysis on Rab8a, leading to its rapid disengagement from TBC1D15L. This rapid turnover potentially facilitates the continuous and premature inactivation of new Rab8a-GTP molecules (Figure 8B), perturbing their trafficking functions.

**Figure 8.**
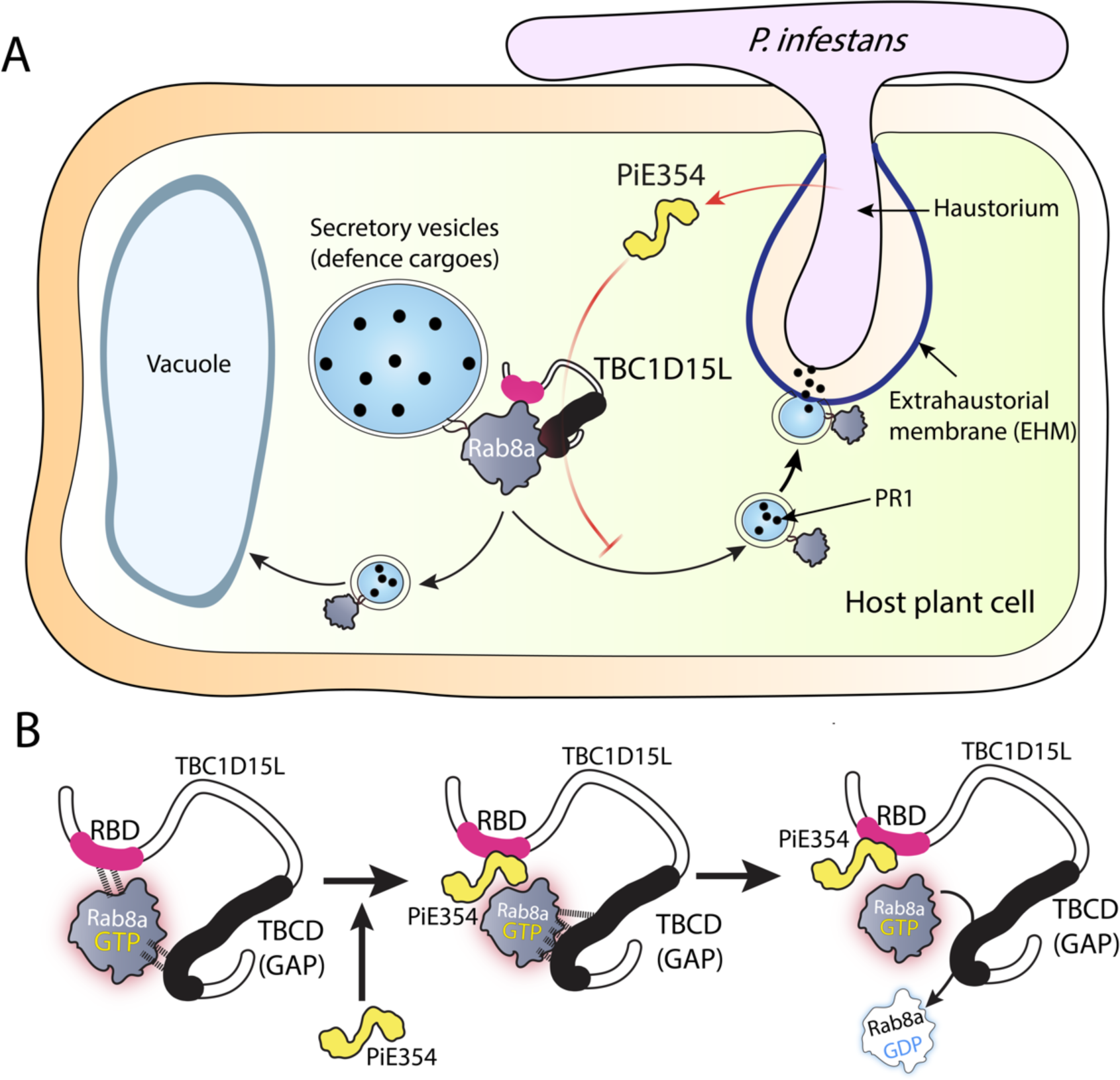
Summary models of the effector PiE354 co-opting the RabGAP TBC1D15L and suppressing plant immunity by redirecting defense-related secretion. (A) Following penetration by *P. infestans* into the host plant cell, the effector PiE354 is secreted into the cytoplasm through its haustoria. Subsequently, PiE354 targets the host RabGAP protein TBC1D15L, facilitating TBC1D15L to deactivate its cognate RabGAP Rab8a. This leads to the redirection of Rab8a trafficking towards the vacuole instead of the plasma membrane, resulting in the suppression of plant immunity by inhibiting antimicrobial PR1 secretion towards the pathogen interface. (B) In the absence of pathogen effectors, Rab8a interacts with both the RBD and TBCD of TBC1D15L. Introduction of the effector PiE354, which binds to the RBD, causes a shift in the positioning of Rab8a towards the TBCD, where the GAP function is housed. Consequently, this facilitates the deactivation of Rab8a through increased GTP hydrolysis.

## Discussion

Accumulating evidence points to host vesicle trafficking as a major hub targeted by pathogen effectors, though the underlying mechanisms are not well understood. Here we uncovered an unprecedented immune subversion mechanism by which *Phytophthora* effectors, PiE354 and its homologs, remodel host membrane dynamics to prevent defense-related secretion through co-opting the host RabGAP protein TBC1D15L. This manipulation effectively redirects defense-related trafficking governed by Rab8a, underscoring the pivotal but previously unknown role of TB1D15L in plant defense. Our findings provide a comprehensive molecular framework illustrating how pathogen effectors co-opt host regulatory components to transform the host-pathogen interface by subverting key immune pathways such as defense-related secretion.

### How does PiE354 co-opt TBC1D15L to subvert Rab8a-mediated trafficking?

A key aspect of our findings is the elucidation of how PiE354 co-opts TBC1D15L to manipulate Rab8a- mediated trafficking. We identified that DUF3548, reported to function as a Rab-binding domain (RBD) in the mammalian TBC-containing RabGAP protein RUTBC2^22^, fulfills a similar role in plant RabGAPs (Figures 5C and 5D). While some RabGAPs like TBC1D15L are characterized by the presence of an RBD, the functional principles of the domain remained enigmatic. Our findings contribute to this understanding by revealing that PiE354 targets the RBD of TBC1D15L, exploiting its GAP function on Rab8a (Figure 7). This association hints at an intramolecular regulatory role for the RBD in modulating the GAP activity.

Building on this, our findings using AF2-M models (Figure 5C) and co-IP experiments (Figure 5D) indicate that Rab8a dynamically interacts with both the Rab-binding and TBC domains of TBC1D15L. We propose that PiE354 binding to the RBD does not necessarily cause a premature release of GTP-bound Rab8a from TBC1D15L, which would otherwise prevent Rab8a inactivation by TBC1D15L. Instead, the docking of PiE354 at the RBD seems to guide Rab8a towards the TBC domain, as indicated by structural predictions (Figures 5C, 7A and 8). This dynamic might negate any restrictive role of the RBD—such as hindering the access of Rab8a to the catalytic GAP interface—thereby boosting the conversion of Rab8a from its active GTP-bound state to its inactive GDP-bound state. However, further studies are needed to comprehensively understand the role of RBDs in TBC-containing RabGAP proteins across various species.

### *Phytophthora* effectors converge on Rab8a-mediated vesicle trafficking pathways

Our research highlights the significance of the Rab8 family of GTPases in plant immunity. This finding resonates with the known roles of the Rab8 family in polarized secretion in both animals and yeast^25^, and its implication in plants for defense-related trafficking of pattern recognition receptors (PRRs) to the plasma membrane^3^. Consistent with the role of Rab8 in polarized secretion in other eukaryotes, our earlier work observed Rab8a-associated vesicles around *P. infestans* haustoria^14^. This study further reveals that Rab8a-mediated trafficking is a key pathway in plant immunity, targeted by effectors from *P. infestans and P. palmivora* that co-opt TBC1D15L, the cognate GAP of Rab8a. This aligns with findings which PexRD54, a *P. infestans* effector, redirects a subset of Rab8a towards autophagy^14^, and RXLR242 from *P. capsici*, which hampers PR1 secretion by interacting with Rab8 members^3^. It is likely that other pathogen effectors have evolved strategies to counter the functions of Rab8a-mediated trafficking, as highlighted by an earlier study indicating the positive role of a Rab8 member in antibacterial immunity^26^. These patterns align with the notion that effectors from the same or different pathogens converge on key immune pathways^27^, emphasizing the critical role of Rab8a in plant defense. Although our study underscores the crucial role of TBC1D15L in regulating Rab8a- mediated trafficking in plant defense, further research is needed to determine if TBC1D15L has regulatory roles across other Rab GTPases.

### PiE354-mediated remodeling of the pathogen interface

The PiE354-mediated subversion of host immunity exemplifies an elaborated strategy employed by pathogen effectors to remodel the host-pathogen interface. By depleting Rab8a from the plasma membrane, PiE354 effectively hampers the secretion of PR1 into the apoplast (Figures 4C and 4D), highlighting a critical pathogen strategy in steering immune components away from the pathogen interface. This mechanism provides insight into the unique biochemistry of membrane interfaces at the pathogen incursion points, such as the formation of haustoria, indicative of a host susceptibility state.

In conclusion, our study illuminates the intricate interplay between plant vesicle trafficking and pathogen effectors, revealing an unprecedented immune evasion strategy to reconfigure pathogen interface. Specifically, we have shown that the effector PiE354 targets the plant RabGAP protein TBC1D15L, effectively subverting defense-related trafficking governed by Rab8a. Our findings thus elucidate a sophisticated evolutionary adaptation by a pathogen effector, which capitalizes on the catalytic functionality of a host transport regulator to compromise innate immune responses.

## Materials and Methods

### Molecular cloning

The molecular cloning of TBC1D15L, TBC1D15L^7M^, TBC1D15L^GAP^, RBDF, TBCDF, PiE354, PiE354^6A^, and PiE355 were conducted using Gibson Assembly, following the methods described in previous works^24,28^. Specifically, the vector backbone is a pK7WGF2 derivative domesticated for Gibson Assembly. The desired sequence for cloning was either manufactured as a synthetic fragment or amplified using designed primers. The fragments were then inserted into the vector using Gibson Assembly, and then transformed into DH5α chemically-competent *E. coli* through heat shock. These plasmids were subsequently amplified and extracted by PureYield™ Plasmid Miniprep System (Promega), and electroporated into *Agrobacterium tumefaciens* GV3101 electrocompetent cells. Sequencing was done by Eurofins. LB agar containing gentamicin and spectinomycin was used to grow constructs carrying pK7WGF2 plasmid. TIKI DNA was synthesized including an N-terminal FLAG tag and flanking attL1, attL2 sites in pUC57, FLAG-linker replaces the signal peptide. For the RNAi interference silencing construct of TBC1D15L (RNAi:TBC1D15L), an intron-containing hairpin RNA vector for RNA interference in plants (pRNAi-GG) was employed, based on Golden Gate cloning method described in a previous study^29^. RNAi:TBC1D15L targeted the region between 1790 and 2031 bp of *TBC1D15L*. The target fragment (TBC1D15L-silencing_synfrag) was synthesized and then inserted into the pRNAi-GG vector both in sense and anti-sense orientation, utilizing the overhangs left by BsaI cleavage. This resulted in the expression of a construct that folds back onto itself, forming the silencing hairpin structure. The subsequent steps of *E. coli* transformation, Miniprep, sequencing, and agrobacterium transformation were the same as those used for the overexpression constructs. LB agar containing gentamicin, kanamycin and chloramphenicol was used to grow constructs carrying pRNAi-GG plasmid. All primers and synthetic fragments used in this study are detailed in **Table S5.** All constructs used in this study are detailed in **Table S6**.

### Plant material

*Nicotiana benthamiana* plants were cultivated in a controlled growth chamber at a temperature of 24°C, using a mixture of organic soil (3:1 ratio of Levington’s F2 with sand and Sinclair’s 2-5 mm vermiculite). The plants were exposed to high light intensity and subjected to a long day condition (16 hours of light and 8 hours of darkness photoperiod). The experiments were performed using plants that were 4 to 5 weeks old.

### *Phytophthora infestans* growth and infection assays

WT and tdTomato-expressing *Pytophthora infestans* 88069 isolates were cultivated on rye sucrose agar (RSA) media in the dark at 18°C for a period of 10 to 15 days before harvesting zoospores^30^. Zoospore solution was obtained by adding cold water at 4°C to the media and then incubating it at 4°C in the dark for 90 minutes. For the infection assay, 10 µl droplets of zoospore solution containing 50000 spores/ml were applied to the abaxial side of agroinfiltrated leaves. The leaves were then kept in a humid environment. Daylight and fluorescent images were captured at 5 to 7 days post-infection, and both lesion sizes and hyphal growth were measured and analyzed using ImageJ.

### Confocal laser scanning microscopy

The confocal microscopy analyses were conducted three days after agroinfiltration. To image the infiltrated leaf tissue, they were excised using a size 4 cork borer, live-mounted on glass slides, and submerged in wells of dH2O using Carolina observation gel (Carolina Biological). The imaging of the abaxial side of the leaf tissue was performed using either a Leica TCS SP8 inverted confocal microscope equipped with a 40x water immersion objective lens or a Leica STELLARIS 5 inverted confocal microscope equipped with a 63x water immersion objective lens. The laser excitations for GFP, RFP, and BFP tags were Argon 488 nm (15%), DPSS 561 nm and Diode 405 nm, respectively. The emission ranges for GFP, RFP, and BFP tags were 495 - 550 nm, 570 - 620 nm, and 402 - 457 nm, respectively. To prevent spectral mixing from different fluorescent tags when imaging samples with multiple tags, sequential scanning between lines was applied. Confocal images, comprising both Z-stack and single plane images, were analyzed using ImageJ.

### Fluorescence microscopy

Fluorescence microscopy was utilized to visualize the hyphal growth of *P. infestans* expressing tdTomato. The imaging setup consisted of a Leica MZ 16 F microscope coupled with the Leica DFC300 FX Digital Color Camera designed for fluorescence imaging. Infected leaf samples were positioned on a petri dish within the microscope imaging area. The imaging filter employed was DsRed, with an excitation range spanning 510 – 560 nm.

### Structural and sequence analyses

The AF2-multimer was employed via a Google Colab subscription, adhering to the set guidelines^31^. With the aid of the "align" command in UCSF ChimeraX (version 1.7), the AF2 predictions were superimposed onto known structures, and the confidence scores of the AF2 predictions were displayed using the local distance difference test (pLDDT) scores on the lDDT-Cα metric^32^. The scoring scale ranged from 0 to 100, with 100 indicating the highest confidence values. For sequence alignment, the MUSCLE algorithm was utilized^33^, and the resulting alignments were visualized and color-coded using ESPript 3.0^34^. Detailed information on the proteins and sequences used for AF2 can be found in **Table S7**.

### Agrobacterium-mediated transient gene expression in *N. benthamiana*

Agrobacterium-mediated transient gene expression was conducted through agroinfiltration, following the previously established method^1^. *Agrobacterium tumefaciens* carrying the desired plasmid was washed with water and then resuspended in agroinfiltration buffer (10 mM MES, 10 mM MgCl2, pH 5.7). The OD600 of the bacterial suspension was measured using the BioPhotometer spectrophotometer (Eppendorf). Subsequently, the suspension was adjusted to the desired OD600 based on the construct and the specific experiment. The adjusted bacterial suspension was then infiltrated into 3 to 4-week-old *N. benthamiana* leaf tissue using a needleless 1ml Plastipak syringe.

### RNA isolation, cDNA synthesis, and RT-PCR

To perform RNA extraction, 56 mg of leaf tissue was promptly frozen in liquid nitrogen. The RNA extraction process employed TRIzol RNA Isolation Reagent (Invitrogen), following the manufacturer’s guidelines. Subsequently, RNA concentration was quantified using NanoDrop Lite Spectrophotometer (Thermo Scientific). 2 mg of the extracted RNA underwent treatment with RQ1 RNase-Free DNAse (Promega) before being used for cDNA synthesis with SuperScript IV Reverse Transcriptase (Invitrogen). The resulting cDNA was then amplified using Phusion High-Fidelity DNA Polymerase (New England Biolabs). GAPDH level was utilized as the transcriptional control. The RT-PCR for *TBC1D15L* was performed using the primers TBC1D15L_RTPCR_F and TBC1D15L_RTPCR_R, while the RT-PCR for *GAPDH* was performed using the primers GAPDH_RTPCR_F and GAPDH_RTPCR_R. All primers used in this study are detailed in **Table S5.**

### Co-immunoprecipitation and immunoblot analyses

Proteins were transiently expressed in *N. benthamiana* leaves through agroinfiltration, and the harvest took place 3 days after agroinfiltration. For western blotting experiments, six leaf discs were excised using a size 4 cork borer (42 mg in total). Meanwhile, co-immunoprecipitation experiments utilized 2 g of leaf tissues. The procedures for protein extraction, purification, and immunoblot analysis followed the previously described protocols^1^. The primary antibodies used included polyclonal anti-GFP produced in rabbit (Chromotek), polyclonal anti-PR1 produced in rabbit (Agrisera), monoclonal anti-RFP produced in mouse (Chromotek), and monoclonal anti-HA produced in rat (Chromotek). As for secondary antibodies, anti-rabbit antibody for HRP detection and AP detection (Sigma-Aldrich), anti-mouse antibody for HRP detection (Sigma-Aldrich), and anti-rat antibody for HRP detection (Sigma-Aldrich) were employed. Comprehensive information regarding the antibodies used is detailed below.

| Primary/Secondary | Detail | Company | Product Code |
| --- | --- | --- | --- |
| Primary | Anti-GFP (Rabbit polyclonal) | Chromotek | PABG1 |
| Primary | Anti-RFP (Mouse monoclonal) | Chromotek | 6G6 |
| Primary | Anti-HA (Rat monoclonal) | Chromotek | 7C9 |
| Primary | Anti-PR1 (Rabbit polyclonal) | Agrisera | AS10 687 |
| Secondary | Anti-rabbit (HRP) | Sigma-Aldrich | A9169 |
| Secondary | Anti-rabbit (AP) | Sigma-Aldrich | SAB3700834 |
| Secondary | Anti-rat (HRP) | Sigma-Aldrich | A9037 |
| Secondary | Anti-mouse (HRP) | Sigma-Aldrich | SAB3700986 |

**List of antibodies used in this research.**

### *Phytophthora infestans* extract preparation and injection

Mycelia obtained from *P. infestans* RSA plates were collected and suspended in 5 ml of water per petri dish. The suspension was vortexed for 1 minute and subsequently heated at 95°C for 20 minutes. Then, the mixture was filtered through filter paper with a 5-13 µm pore size. The resulting filtrate underwent an additional filtration step using a syringe filter with a 0.45 µm pore size. This resultant solution was then administered to plants to serve as a PAMP cocktail.

### Apoplastic washing fluid extraction

Apoplastic proteins were obtained following the previously described procedure^35^. Detached and washed *N. benthamiana* leaves, which had undergone infiltration, were rolled up and placed into a needleless syringe containing distilled water. Negative pressure was generated within the syringe to facilitate the infiltration of the entire leaves with water. Afterwards, the infiltrated leaves were wrapped in parafilm, placed into a syringe with the plunger removed, and inserted into a Falcon tube. The entire setup was then centrifuged for 10 minutes at 1000 g at 4°C. The apoplastic washing fluid accumulated at the bottom of the tube was collected and promptly frozen using liquid nitrogen. The remaining leaf tissue was gathered for subsequent immunoblotting analysis.

### Cell death assay

Cell death elicitors were introduced into the abaxial side of *N. benthamiana* leaves through agroinfiltration. Subsequently, at 2-4 dpi, the leaves were detached and subjected to imaging under both daylight and UV light conditions. The intensity of cell death was evaluated using a well-established seven-tiered cell death index^36^.

### Rab8a purification

His-tagged Rab8a in Rosetta2 (DE3) pLysS strain *E. coli* was received from Yasin Dagdas. In brief, *E. coli* transformed with Rab8a was grown to OD600 of 0.6 and induced overnight with 0.3 mM IPTG at 18°C. Harvested cells were frozen in -80°C until needed. Cells were resuspended in 100 mM sodium phosphate pH 7.2, 300mM NaCl, 50mM imidazole (buffer A) with an EDTA-free protease inhibitor tablet (cOmplete^TM^, Roche). Following cell disruption (CF Cell Disrupter (Constant Systems Ltd) at 27 kpsi, 4°C, for 3 times) and ultracentrifugation, clear lysate was applied to a 5 mL HisTrap HP (Cytiva) using a peristaltic pump (P1, Cytiva). Elutions were performed using an imidazole gradient (buffer A and buffer A + 500mM imidazole) applied by an ÄKTA pure protein purification system (Cytiva). Fractions were analysed with SDS-PAGE and eluted Rab8a was pooled and concentrated using an Amicon Ultra 30kDa centrifugation filter. Rab8a was dialysed into 100 mM sodium phosphate pH 7.2, 150 mM NaCl buffer and stored at -80°C. Using purified Rab8a above the concentration of 0.1 mg/ml led to a reduction in GTP levels, verifying the functionality of the purified Rab8a (Figure S5E). We decide to carry out subsequent GTPase activity assays with 0.2 mg/ml Rab8a since this concentration gave a robust and clear GTPase readout.

### GTPase activity assay

To investigate the impact of proteins of interest on the GTPase activity of Rab8a, we employed a luciferase-based GTPase assay utilizing the GTPase-Glo^TM^ Assay Kit (Promega). The assay was conducted following the guidelines provided by the manufacturer. Specifically, a mastermix of 2X GTP-GAP solution was prepared, containing 10 μM GTP and 1 mM DTT in GTPase/GAP buffer. In each well of the microplate, 12.5 μl of 0.4 mg/ml Rab8a was added, which was diluted in the buffer provided. The GTPase reaction was initiated by adding 12.5 μl of the 2X GTP-GAP solution to each well. The reaction was incubated for 120 minutes at RT with continuous shaking. Upon completion of the GTPase reaction, 25 μl of reconstituted GTPase-Glo^TM^ Reagent was introduced to convert the unhydrolyzed GTP to ATP. The plate was then incubated for 30 minutes at RT with shaking. Subsequently, 50 μl of Detection Reagent was added to all the wells, and they were incubated for 10 minutes at RT. Finally, luminescence was measured using CLARIOstar® Plus plate reader.

### Image processing and data analysis

The confocal microscopy images were processed using Leica LAS X software and ImageJ. Depending on the specific experiment, the confocal images could be either single plane images or Z-stack images, and this information is provided in the figure legends. Image analysis and quantification for cell death

and infection assay experiments were performed using ImageJ. For data representation, violin plots and box plots were created using R, while bar graphs were generated using Microsoft Excel. To assess statistical differences, a range of tests, including Student’s t-test and Mann-Whitney U test, were conducted based on statistical normality and variance. Measurements were deemed significant when p < 0.05 (*), p < 0.01 (**) and highly significant when p < 0.001 (***). Detailed information regarding all the statistical calculations can be found in **Table S8**.

### Accession numbers/identifiers

TBC1D15L (Nbe.v1.s00100g29830) TIKI (PLTG_0964243, Table S1) PiE354 (PITG_04354, Table S1) PiE355 (PITG_04355, Table S1)

## Acknowledgments

We extend our sincere appreciation to the Imperial College FILM facility for their technical expertise and provision of microscopy equipment. We would also like to acknowledge Mehdi Doumane and Ayoub Kadoussi for their technical contributions. Additionally, we would like to express our gratitude to the members of the Bozkurt, Schornack, and Kamoun groups for their insightful suggestions and engaging discussions. E.L.H.Y. would like to thank Baptiste Castel, Cécile Segonzac, Mark Banfield, Şuayib Üstün, and Thorsten Langner for providing useful feedback and encouragement during the OMGN 2022, MPMI 2023 and ICPP 2023 conferences.

## Funding

BBSRC grant BB/X016382/1 (E.L.H.Y.) BBSRC grant BB/T006102/1 (Y.T., T.O.B.)

BBSRC Impact Acceleration Fund BB/X511055/1 (T.I.)

## Author contributions

Conceptualization: E.L.H.Y., S.K., S.S., T.O.B.

Methodology: E.L.H.Y., S.S., T.O.B., E.E., F.T.,

Validation: E.L.H.Y., Y.T., L.I.C., E.E., F.T.,

Formal Analysis: E.L.H.Y., Y.T., L.I.C., T.I., E.E., F.T., J.S., F.M.

Investigation: E.L.H.Y., Y.T., L.I.C., T.I., E.E., F.T., J.S., F.M.

Data Curation: E.L.H.Y., S.S., T.O.B.

Visualization: E.L.H.Y., T.O.B.

Writing – Original Draft: E.L.H.Y., T.O.B. Writing – Review & Editing: E.L.H.Y., T.O.B.

Supervision: E.L.H.Y., F.M., S.K., D.B., S.S., T.O.B.

Funding Acquisition: S.K., D.B., S.S., T.O.B.

## Competing interest statement

T.O.B. and S.K. receive funding from industry on NLR biology. T.O.B. and S.K. cofounded Resurrect Bio Ltd. on resurrecting disease resistance. The remaining authors have no conflicts of interest to declare.

## Declaration of generative AI and AI-assisted technologies in the writing process

During the preparation of this work, the authors used ChatGPT to check grammar and phrasing. After using ChatGPT, the authors reviewed and edited the content as needed and take full responsibility for the content of the publication.

## Data and materials availability

All relevant study data are included in the article, and in the Supplementary Materials files. AF2- multimer predictions are uploaded to the public repository Figshare and is available at https://doi.org/10.6084/m9.figshare.24846558.

## Supplementary materials in this combined PDF include

Supplementary figures s1-S6

## Other supplementary materials for this manuscript include

Table S1. Protein and DNA sequences of effectors Table S2. List of interactors of TIKI identified in Y2H Table S3. List of interactors of TIKI identified by IP-MS

Table S4. List of interactors of TBC1D15L identified by IP-MS Table S5. Primers and synthetic fragments used

Table S6. Detail of all constructs

Table S7. Proteins and sequences used for AF2 Table S8. Statistics details and summary

---

**Figure S1.**
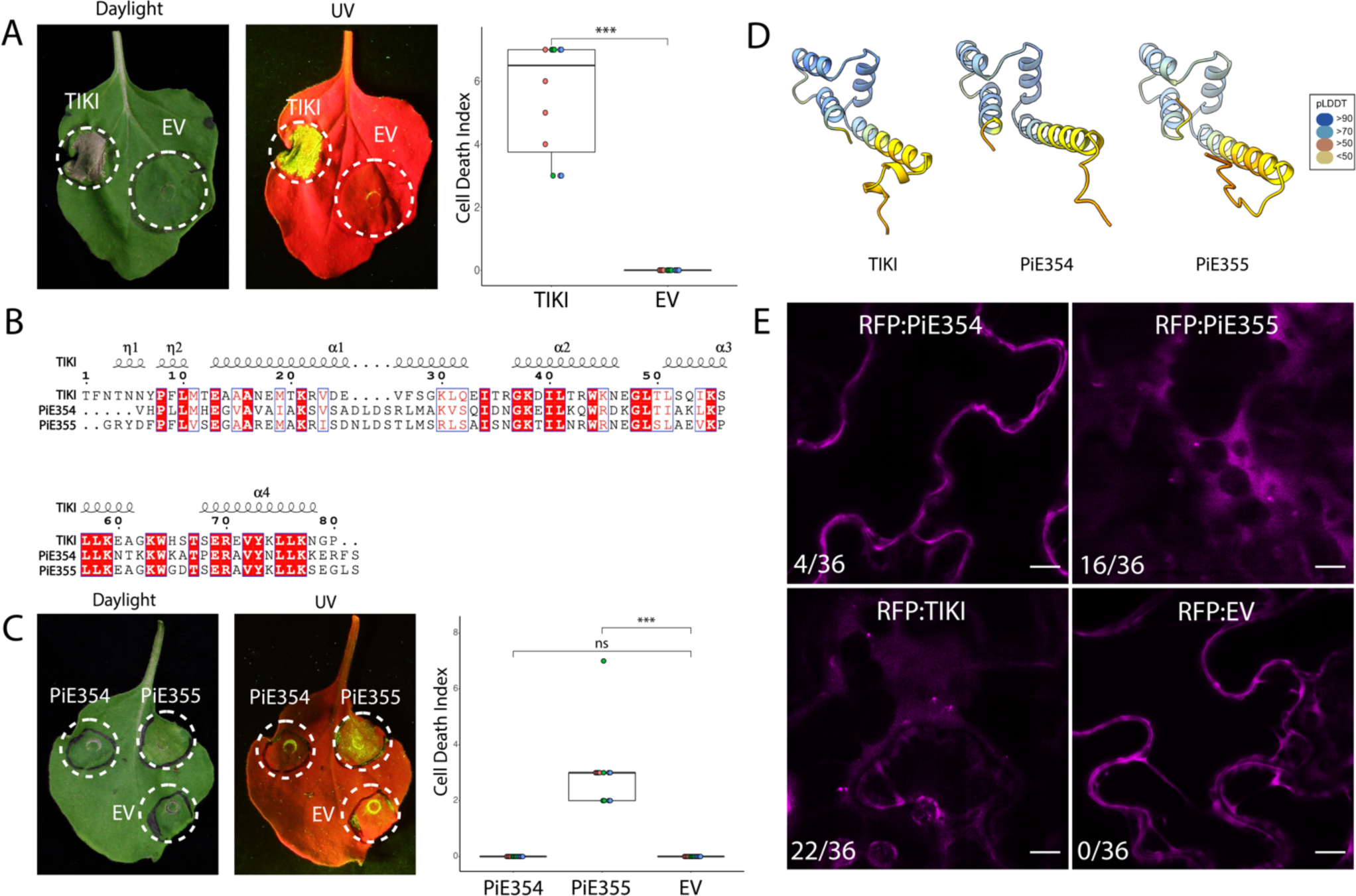
Conserved effectors from *Phytophthora* species target TBC1D15L. Related to Figure 1. (A) Expression of TIKI causes cell death. Representative *N. benthamiana* leaves infiltrated with TIKI or EV. Daylight and UV images were taken at 4 dpi, and cell death was scored at 4 dpi. Box and dot plot showing TIKI expression causes cell death in plants (5.5, N = 12), while EV control does not (0.0, N = 12). (B) Pairwise amino acid sequence alignment of the *Phytophthora* effectors TIKI, PiE354 and PiE355. Alignments were obtained using the MUSCLE algorithm and were visualized and color-coded via ESPript 3.0^34^. The squiggle symbols on top indicate α-helices. Red box indicates strict identical residues, while blue frame indicates similar residues. (C) Expression of PiE355, but not PiE354, causes cell death. Representative *N. benthamiana* leaves infiltrated with PiE354, PiE355, or EV. Daylight and UV images were taken at 4 dpi, and cell death was scored at 4 dpi. Box and dot plot showing PiE355 expression causes cell death in plants (3.0, N = 12), while PiE354 expression (0.0, N = 12) and EV expression (0.0, N = 12) do not. (D) AF2 structures of the effectors TIKI from *P. palmivora*, and PiE354 & PiE355 from *P. infestans*. The colors of TIKI, PiE354 and PiE355 are based on the AF2-calculated prediction confidence score (pLDDT) as indicated in the rectangular box. (E) PiE354 forms punctate structures less frequently than PiE355 and TIKI. Confocal micrographs of *N. benthamiana* leaf epidermal cells transiently expressing RFP:PiE354, RFP:PiE355, RFP:TIKI, or RFP:EV control. For RFP:PiE354, 4 out of 36 images contain punctate structures. For RFP:PiE355, 16 out of 36 images contain punctate structures. For RFP:TIKI, 22 out of 36 images contain punctate structures. For RFP:EV, none of the 36 images contain punctate structures. Presented images are single plane images. Scale bars, 5 µm. All statistical differences were analyzed by Mann-Whitney U test in R. Measurements were highly significant when p < 0.001 (***).

**Figure S2.**
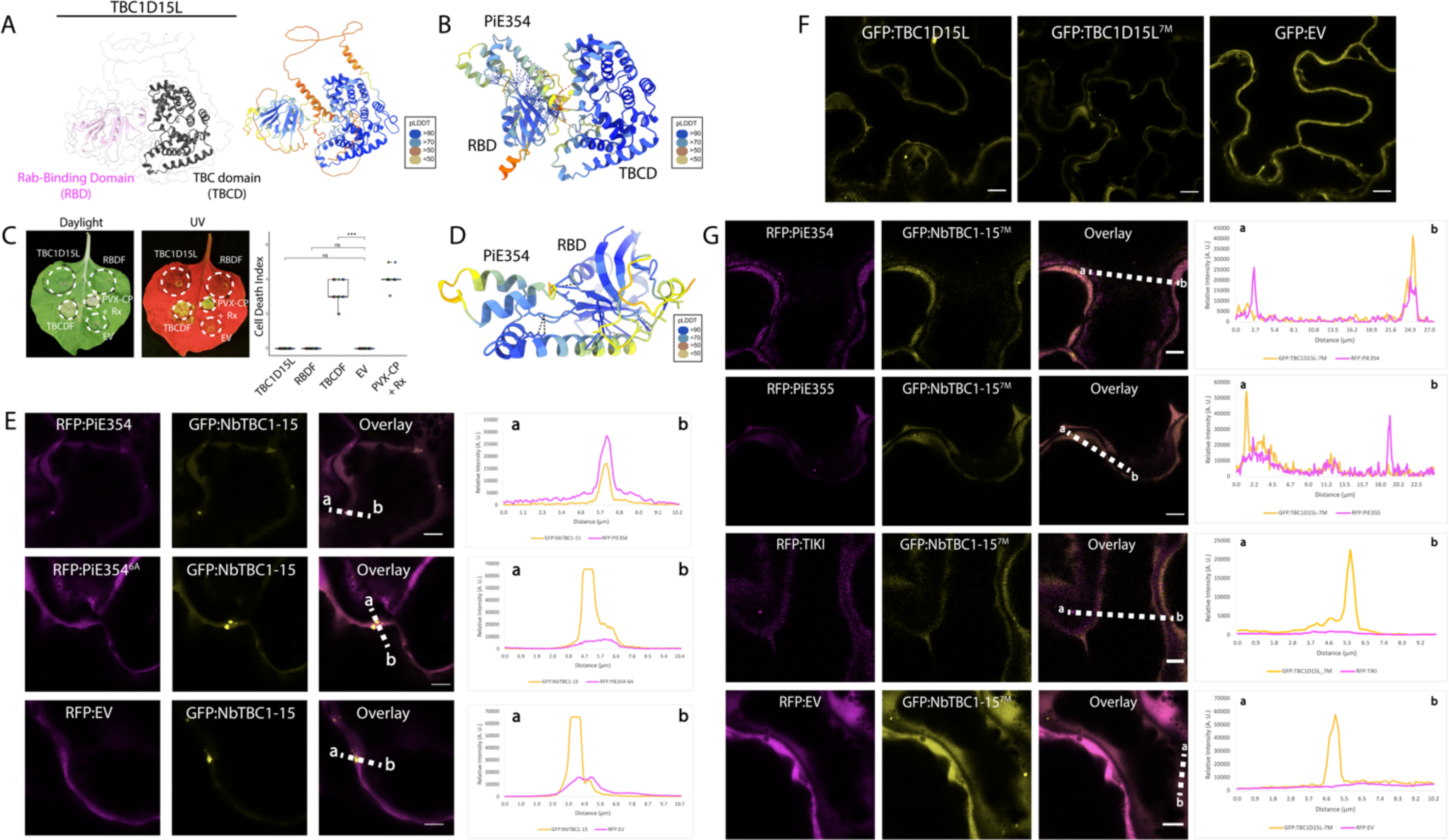
PiE354 targets the N-terminal RBD fragment of TBC1D15L. Related to. Figure 2. (A) AF2 model of TBC1D15L architecture. (Left panel) A Rab-binding domain (RBD) is located at the N-terminal, and a TBC domain (TBCD) is located at the C-terminal. (Right panel) The colors of the AF2 model of TBC1D15L are based on the AF2-calculated prediction confidence score (pLDDT) as indicated in the rectangular box. (B) AF2-M-predicted model of PiE354 targeting TBC1D15L. The colors of the PiE354-TBC1D15L AF2-M model are based on the AF2-calculated prediction confidence score (pLDDT) as indicated in the rectangular box. (C) Expression of TBCDF elicits cell death activity. Representative *N. benthamiana* leaves infiltrated with TBC1D15L, RBDF, TBCDF or EV control. PVX-CP and Rx were co-infiltrated as a positive control for cell death activity. Daylight and UV images were taken at 4 dpi, and cell death was scored at 4dpi. Box and dot plot showing TBCDF expression causes cell death in plants (3.3, N = 20) while TBC1D15L (0.0, N = 20), RBDF (0.0, N = 20) and EV control expression (0.0, N = 20) do not. Statistical differences were analyzed by Mann-Whitney U test in R. Measurements were highly significant when p<0.001 (***). (D) AF2-M-predicted model of PiE354 targeting the RBD of TBC1D15L. The colors of the PiE354-RBD AF2-M model are based on the AF2- calculated prediction confidence score (pLDDT) as indicated in the rectangular box. (E) PiE354 colocalizes with TBC1D15L through 6 key residues on PiE354. Confocal micrographs of *N. benthamiana* leaf epidermal cells transiently expressing either RFP:PiE354 (1^st^ row), RFP:PiE354^6A^ (2^nd^ row), or RFP:EV (3^rd^ row), with GFP:TBC1D15L. (F) TBC1D15L and TBC1D15L^7M^ show cytoplasmic localization with puncta formation. Confocal micrographs of *N. benthamiana* leaf epidermal cells transiently expressing GFP:TBC1D15L, GFP:TBC1D15L^7M^, or GFP:EV control. (G) TBC1D15L^7M^ does not colocalize with the effectors PiE354, PiE355 and TIKI in puncta. Confocal micrographs of *N. benthamiana* leaf epidermal cells transiently co-expressing GFP:TBC1D15L^7M^ with either RFP:PiE354 (1st row), RFP:PiE355 (2nd row), RFP:TIKI (3rd row), or RFP:EV control (4th row). All presented confocal images are single plane images. Transects in overlay panels correspond to line intensity plots depicting the relative fluorescence across the marked distance. Scale bars, 5 µm.

**Figure S3.**
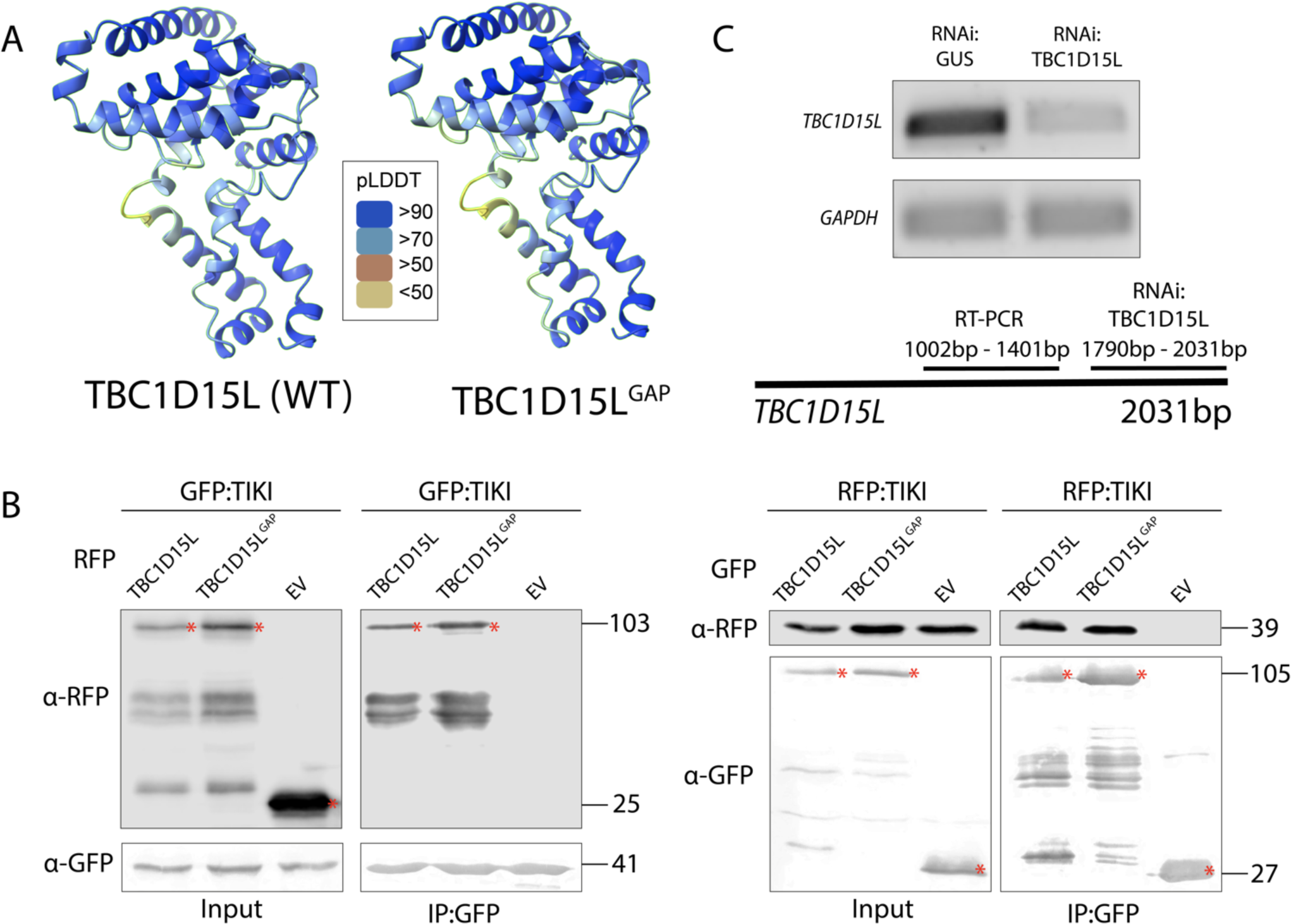
TBC1D15L negatively regulates plant immunity through its GAP function. Related to. Figure 3. (A) AF2-predicted structures of TBC1D15L and its GAP mutant TBC1D15L^GAP^. The colors of TBC1D15L and TBC1D15L^GAP^ are based on the AF2-calculated prediction confidence score (pLDDT) as indicated in the rectangular box. (B) TIKI interacts with TBC1D15L *in planta* independent of the GAP function of TBC1D15L. (Left panel) GFP:TIKI was transiently co-expressed with either RFP:TBC1D15L, RFP:TBC1D15L^GAP^, or RFP:EV. IPs were obtained with anti-GFP antibody. (Right panel) RFP:TIKI was transiently co-expressed with either GFP:TBC1D15L, GFP:TBC1D15L^GAP^, or GFP:EV. IPs were obtained with anti-GFP antibody. Total protein extracts were immunoblotted. Red asterisks indicate expected band sizes. Numbers on the right indicate kDa values. (C) Validation of *TBC1D15L* silencing by RNAi:TBC1D15L. Constructs carrying hairpin plasmids (pRNAi-GG) targeting TBC1D15L or the GUS reporter gene were infiltrated to *N. benthamiana* leaves. The expression levels of the targeted genes were assessed via RT-PCR at 3 days post silencing. RT-PCR, employing primers TBC1D15L_RTPCR_F and TBC1D15L_RTPCR_R, confirmed efficient gene silencing of TBC1D15L using the RNAi:TBC1D15L construct. Glyceraldehyde 3-phosphate dehydrogenase (GAPDH) served as the internal control, using primers GAPDH_RTPCR_F and GAPDH_RTPCR_R for assessment. The cDNA was synthesized using total RNA.

**Figure S4.**
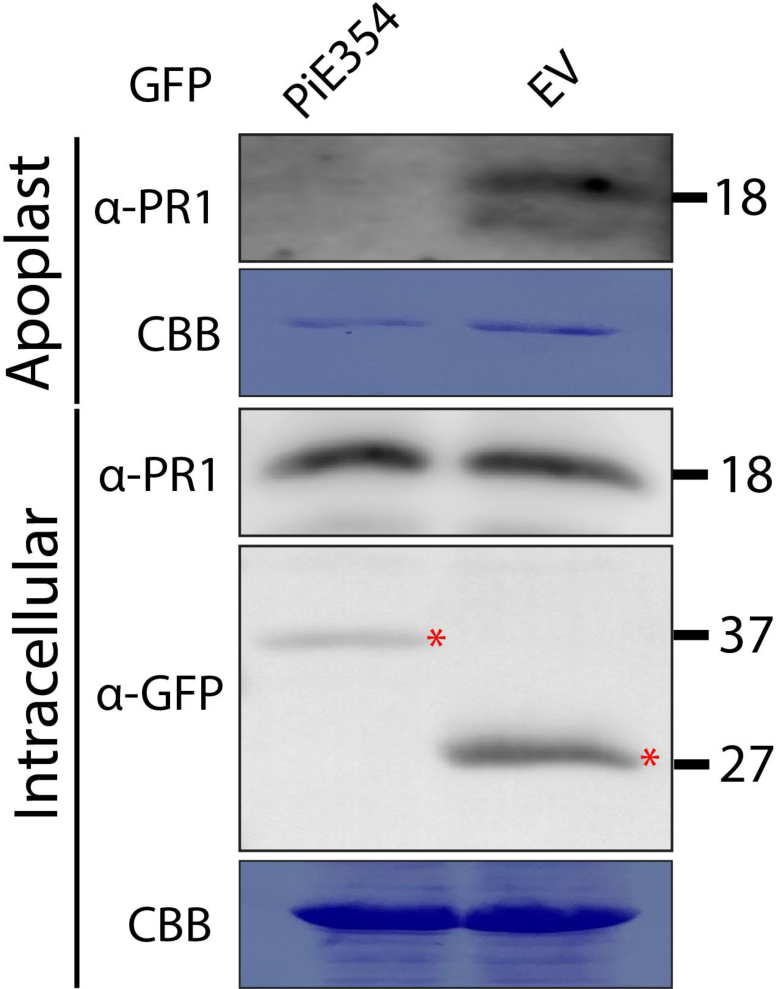
PiE354 disrupts antimicrobial PR1 secretion into the apoplast. Related to. Figure 4. To conduct PR1 secretion assays, the infiltrated leaves were challenged with *P. infestans* extract at 3 dpi and proteins were extracted from the apoplast and leaf tissue at 4 dpi and immunoblotted. Western blot shows PiE354 disrupts antimicrobial PR1 secretion into the apoplast. *N. benthamiana* leaves were infiltrated to express GFP:PiE354, or GFP:EV. Red asterisks show expected band sizes. Numbers on the right indicate kDa values.

**Figure S5.**
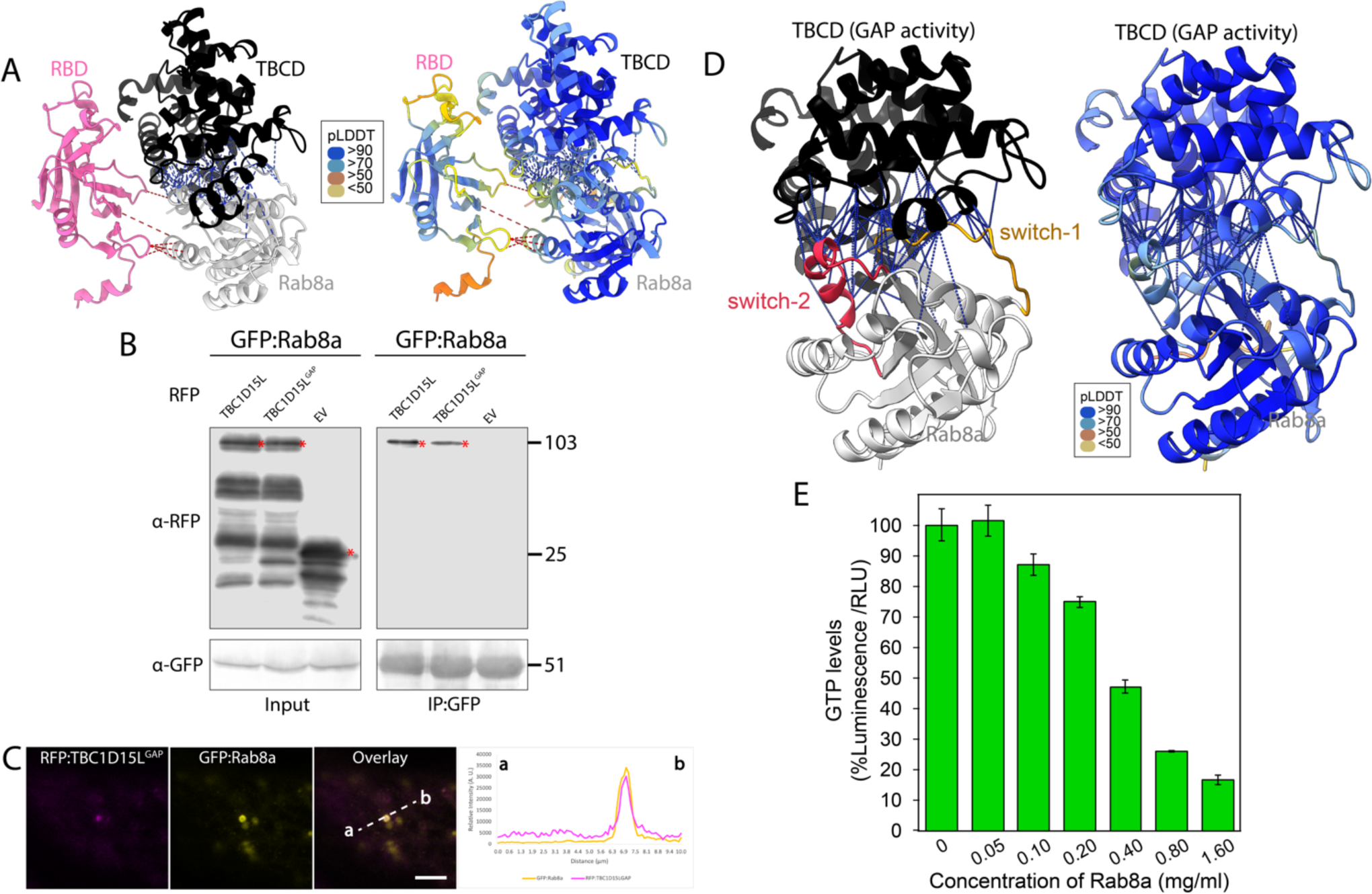
Rab8a is a GAP substrate of TBC1D15L. Related to. Figure 5. (A) AF2-M-predicted model of full length TBC1D15L and Rab8a in complex. (Left panel) Rab8a interacts with both the RBD fragment (RBDF) and TBCD fragment (TBCDF) of TBC1D15L. (Right panel) The colors of full length TBC1D15L-Rab8a AF2-M model are based on the AF2-calculated prediction confidence score (pLDDT) as indicated in the rectangular box. (B) Rab8a interacts with TBC1D15L *in planta* independent of the GAP activity of TBC1D15L. GFP:Rab8a was transiently co-expressed with either RFP:TBC1D15L, RFP:TBC1D15L^GAP^, or RFP:EV. IPs were obtained with anti-GFP antibody. Total protein extracts were immunoblotted. Red asterisks indicate expected band sizes. Numbers on the right indicate kDa values. (C) Rab8a colocalizes with TBC1D15L in puncta independent of the GAP activity of TBC1D15L. Confocal micrographs of *N. benthamiana* leaf epidermal cells transiently expressing RFP:TBC1D15L^GAP^ with GFP:Rab8a. Presented images are single plane images. Transects in overlay panels correspond to line intensity plots depicting the relative fluorescence across the marked distance. Scale bars, 5 µm. (D) AF2-M-predicted model of Rab8a in a complex with the TBCD of TBC1D15L. (Left panel) TBCD, which is crucial for the GAP activity of TBC1D15L, makes multiple contacts with the switch-1 and switch-2 regions of Rab8a that regulate GTP hydrolysis activity. Switch-1 and switch-2 regions that are flanking the GTP binding pocket are colored bronze and red respectively. (Right panel) The colors of TBCDF-Rab8a AF2-M model are based on the AF2- calculated prediction confidence score (pLDDT) as indicated in the rectangular box. (E) Bar graph depicting the impact of varying concentrations of Rab8a on intrinsic GTPase activity. Serially diluted Rab8a in GTPase/GAP Buffer was combined with 2X GTP solution containing 10 μM GTP and 1 mM DTT, resulting in initial reaction mixtures with Rab8a concentrations ranging from 0 to 1.60 mg/ml. These mixtures were then incubated at 25°C for 120 minutes as per manufacturer’s guidelines. An equal volume of reconstituted GTPase-Glo™ Reagent was added to the solutions to cease the GTP hydrolysis reaction, and the mixture was incubated for 30 minutes at room temperature. Following the incubation, the detection buffer was added, and luminescence levels were recorded, which correspond to the quantity of unhydrolyzed GTP remaining in the solution after the GTPase reaction.

**Figure S6.**
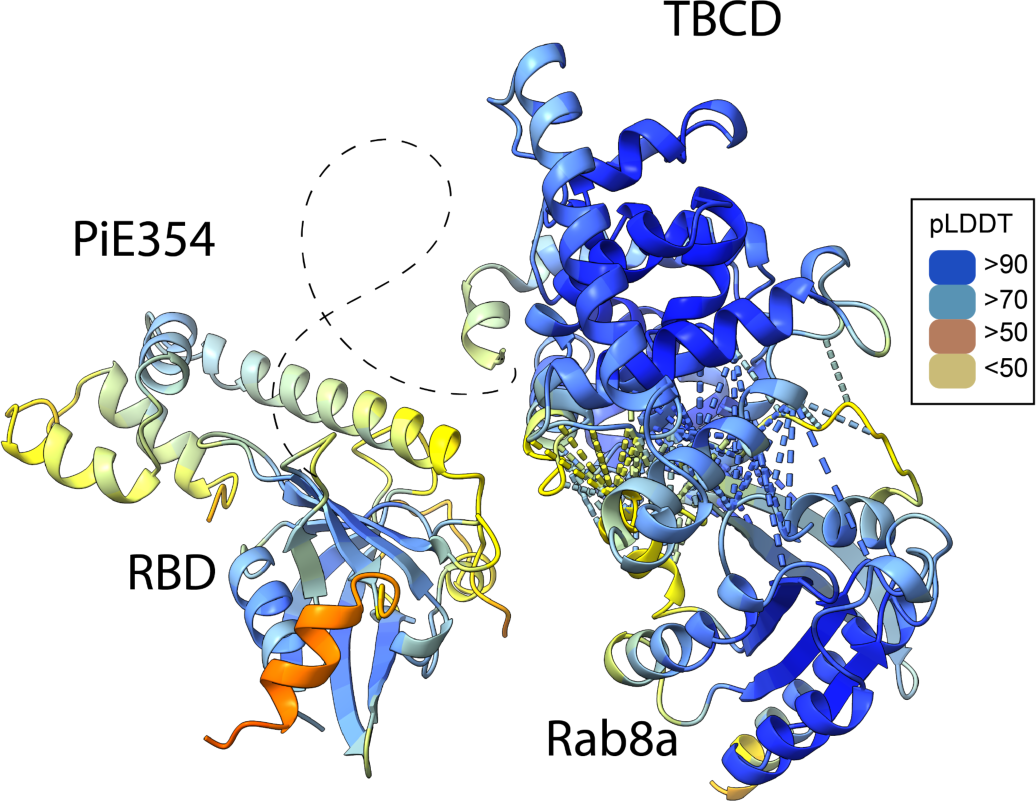
AF2-M-predicted model of PiE354 in complex with the TBC1D15L-Rab8a pair. Related to. Figure 7. The colors of PiE354-TBC1D15L-Rab8a AF2-M model are based on the AF2- calculated prediction confidence score (pLDDT) as indicated in the rectangular box.

